# Gut-derived serine protease reduces sleep but extends lifespan

**DOI:** 10.64898/2026.06.25.734557

**Authors:** Rebecca S. Moore, Richard Xiong, Erick Astacio, Julie Williams, Thomas G. Brooks, Gregory Grant, Joseph Stucynski, Hossein Fazalinia, Erik Nash, Lynn Spruce, Amita Sehgal

**Affiliations:** Department of Neuroscience, Chronobiology and Sleep Institute University of Pennsylvania, Philadelphia, PA, USA; Institute for Translational Medicine and Therapeutics, University of Pennsylvania, Philadelphia, PA, USA; Department of Genetics, University of Pennsylvania, Philadelphia, PA, USA; Proteomics Core Facility, Children’s Hospital of Philadelphia, Philadelphia, PA, USA; Department of Biomedical and Health Information, Children’s Hospital of Philadelphia, Philadelphia, PA, USA; Howard Hughes Medical Institute, University of Pennsylvania, Philadelphia, PA, USA

## Abstract

Sleep has historically been viewed through a brain-centric lens with little known about the contribution of the periphery. Through a targeted screen of secreted peptides from the fat body, gut, and body wall muscle in *Drosophila*, we identified CG11037, a trypsin-like serine endopeptidase secreted from midgut enterocytes, as a previously unrecognized regulator of sleep. Loss of *CG11037* reduces daily sleep, as well as sleep following injury/infection, although recovery is enhanced suggesting reduced need for sleep during sickness. Proteomic analysis of flies with reduced *CG11037* revealed altered oxidative stress response pathways including cytochrome P450–related proteins. Indeed, gut-specific knockdown of *Cyp6a2* or *Cyp28d1* phenocopies *CG11037* sleep defects, and loss of *CG11037* or P450 elevates reactive oxygen species (ROS) selectively in the gut without affecting brain ROS. Rescue of sleep by antioxidant treatment demonstrates that peripheral ROS accumulation drives the behavioral phenotype. Strikingly, despite physiological dysfunction and reduced sleep, *CG11037* or Cytochrome P450 knockdown extends lifespan in a ROS-dependent manner, suggesting that stress adaptation of these knockdowns allows them to live longer. Together, these findings uncover a mechanism for the unexpected association of reduced sleep and extended lifespan.

## Introduction

Sleep is one of the most conserved behavioral states across animals, yet its biological purpose and regulation remain incompletely understood. Sleep research has been profoundly shaped by the view that sleep is fundamentally a brain-generated process. As famously stated by J. Allan Hobson, “sleep is of the brain, by the brain and for the brain” (1). This perspective was influenced by decades of research identifying neural circuits and neurotransmitter systems that regulate sleep and wakefulness (2). However, this brain-centric view is increasingly difficult to reconcile with evidence that sleep is tightly coupled to whole-organism physiology.

Sleep is highly sensitive to metabolic state, immune activation, infection, and cellular stress. Circulating cytokines (3,4), metabolic hormones (5), and stress signals (6,7) can alter sleep duration and architecture suggesting that sleep is not solely regulated within the brain. Overall, these findings challenge the notion that sleep is exclusively a neural process and suggest that sleep regulation requires continuous communication between the brain and peripheral tissues. Despite this growing appreciation for systemic influences, however, the underlying molecular mechanisms remain poorly understood.

The fruit fly *Drosophila melanogaster* provides a powerful model for identifying such mechanisms (8,9). In addition to its well-characterized neural sleep circuits, Drosophila offers genetic tools that allow tissue-specific manipulation of signaling molecules originating from peripheral organs (10). Indeed, emerging evidence implicates molecules produced in the Drosophila periphery in sleep (11–14).

Here, we conducted an unbiased RNAi screen to systematically identify secreted peptides from the fly intestine, fat body, and body wall muscle that regulate daily sleep. We identified *CG11037*, a gut expressed trypsin-like serine endopeptidase, that is required during adulthood for normal daytime and nighttime sleep in males and females. Mechanistically, flies deficient in *CG11037* have reduced expression of several oxidative stress response proteins including Cytochrome P450s suggesting that loss of *CG11037* impairs redox metabolism. Accordingly, CG11037 gut-knockdown elevates baseline levels of reactive oxygen species (ROS) which is required for its’ reduced sleep. Furthermore, gut-specific knockdown of Cytochrome P450 enzymes *Cyp6a2* and *Cyp28d1* phenocopy *CG11037* knockdown. Finally, despite reducing sleep, loss of *CG11037* extends lifespan, which is also dependent on ROS accumulation. Thus, identification of this gut-derived protease provides a mechanism for increased longevity in the face of decreased sleep.

## Results

### RNAi knockdown screen to identify peripheral regulators of sleep

To identify secreted peptides expressed in the gut, fat body, or body wall muscle that regulate daily sleep, we generated a candidate list using a bioinformatic pipeline. We applied the following criteria to develop this list: (1) the gene must encode a protein containing a signal peptide sequence, but no transmembrane domain; (2) the gene must be expressed in the fat body, gut, or muscle(15); (3) the gene must not be expressed in the brain, based on published data for 3 or 20 days post eclosion brains (16). Using these criteria, we identified 388 candidate genes. We then excluded genes with gene ontology annotations associated with embryonic or early developmental processes, reducing the list to 345 genes. Finally, we limited our analysis to genes for which a specific RNAi line was available(17). In total we screened 321 RNAi lines representing 217 genes: 55 uniquely expressed in the digestive tract, 58 uniquely expressed in the fat body, 70 uniquely expressed in the muscle, 3 shared between muscle and digestive tract, and 15 shared between digestive tract and fat body (Table S1). Each gene was selectively knocked down during adulthood in the tissue where it is most highly expressed (Table S1), using inducible (gene-switch) drivers for the gut (TiGS-2) (18), fat body (S106), or muscle (43641). Sleep was measured in 4-6-day-old male and female flies using Drosophila Activity Monitors (19). Flies expressing RNAi targeting candidate genes in the tissue of interest were compared to control flies expressing RNAi against mCherry or GFP in the same tissue.

We identified both sex-specific and shared hits that increased or decreased sleep relative to controls across all tissues examined (Figure 1A). To refine this list, we focused on genes that: (1) affected sleep in both males and females, (2) altered total sleep amount rather than just the distribution of sleep, (3) produced an overall sleep change greater than two standard deviations from the mean control total sleep, and (4) produced a sleep phenotype replicated in subsequent testing. Using these criteria, we identified four hits; two genes whose knockdown in the gut reduced daily sleep, two genes whose knockdown in the fat body reduced daily sleep, and none in the muscle that met these criteria (Figure 1A yellow bars, Table S2). Notably, half of the four final hits are annotated as having serine endopeptidase (serine protease) activity, specifically within the trypsin or chymotrypsin families. Serine endopeptidases are highly prevalent and conserved enzymes, compromising approximately 3% of the *Drosophila* proteome (20). Despite their abundance, their role in sleep regulation has not been previously described.

**Figure 1:**
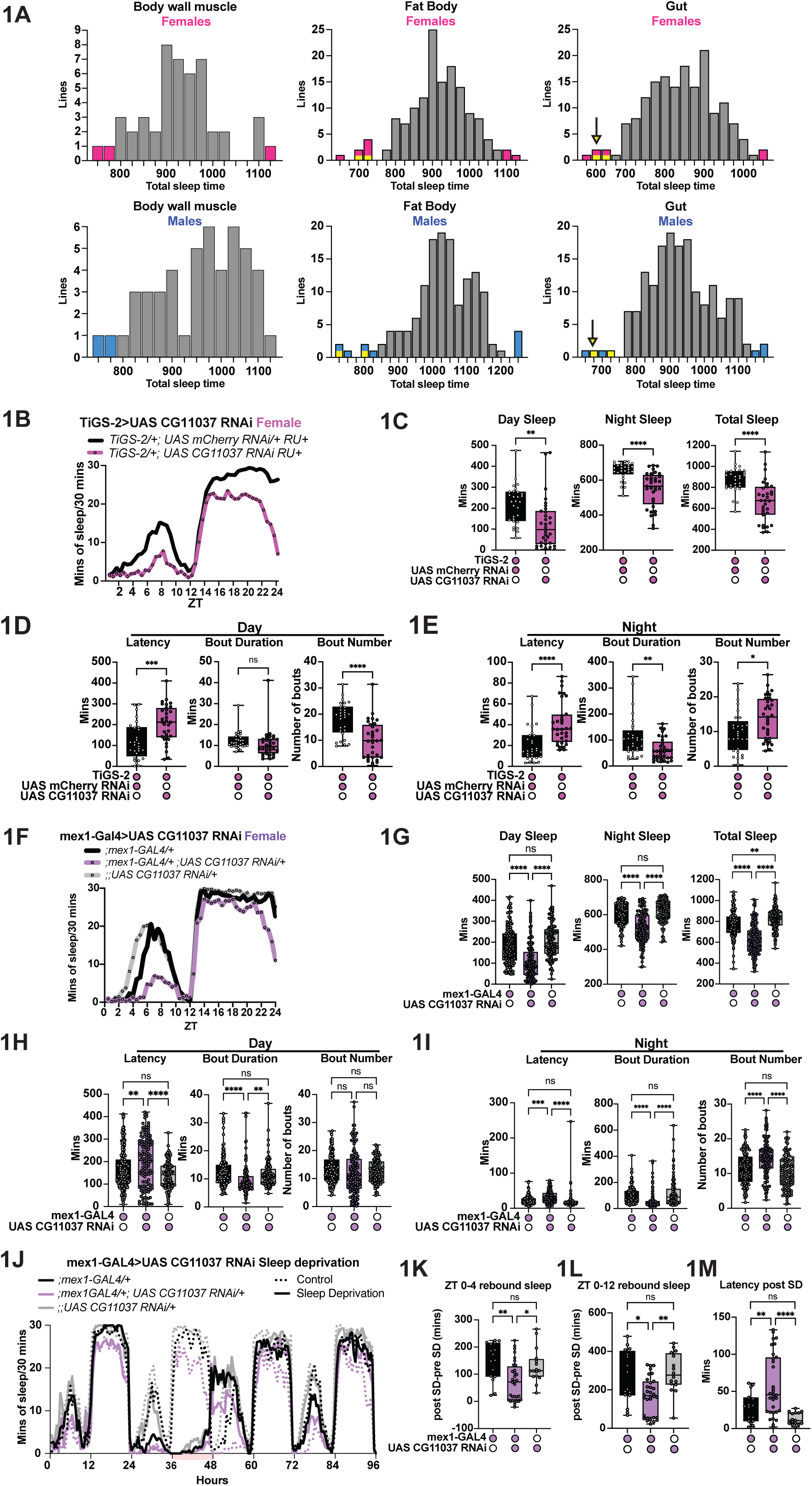
Gut enterocyte secreted *CG11037* regulates daily sleep. (A) Histogram displaying the distribution of sleep over 24 hrs from muscle, fat body, or gut loss-of-function RNAi screen. A *UAS-RNAi* (321 total lines) for each candidate gene (217 total genes tested) was driven by either *MCH-GS* (body wall muscle), *S106-GS* (Fat body), or *TiGS-2* (Gut) by adding RU486 to the food only during sleep recordings. Daily sleep is depicted as the average total sleep time (minutes per 24 hours; x axis) for females (top) and males (bottom). RNAi knockdowns resulting in sleep differences ± two standard deviations from the average control sleep time are indicated in pink or blue for females or males, respectively. Hits that affected sleep in both males and females and were considered for future study are indicated in yellow. An arrow indicates the hit *CG11037*. (B) Representative sleep trace of ;*TiGS-2/+; UAS CG11037 RNAi/+* (pink), and *TiGS-2/+; UAS mCherry RNAi/+* (black) females over the 24-hour day (ZT = Zeitgeber time ZT0-12 = daytime, ZT12-24 = nighttime). (C) Adult gut knockdown of *CG11037* (pink bars) reduces day sleep (left), night sleep (middle), and total sleep (right) in females compared to ;*TiGS-2/+; UAS mCherry RNAi/+* controls (black bars). (D) Adult only gut knockdown of CG11037 (pink) increases daytime sleep latency (left), reduces daytime sleep bouts (right) without affecting daytime bout length (middle) compared to controls (black bars). (E) Similar to daytime sleep, adult gut knockdown of *CG11037* (pink) increases nighttime sleep latency (left). However knockdown flies have more nighttime sleep bouts (right) that are shorter in length (middle) compared to controls (black). (F) Representative sleep trace of *;mex1-GAL4/+; UAS CG11037 RNAi/+* (purple), *GAL4* (black) and *UAS-RNAi* (gray) control females. (G) Whole life enterocyte knockdown of *CG11037* (*mex1-GAL4*) reduces daytime (left), nighttime (middle), and total sleep (right). (H) Constitutive *CG11037* gut knockdown increases daytime latency (left), does not affect bout number (right), but shortens bout duration (middle) compared to *GAL4* and *UAS RNAi* controls. (I) *;mex1-GAL4/+; UAS CG11037 RNAi/+* increases nighttime latency and number of sleep bouts, while bout duration is decreased compared to GAL4 and *UAS-RNAi* controls. (J) Sleep time course of *;mex1-GAL4/+; UAS CG11037 RNAi/+* (purple), *GAL4* (black), and *UAS-RNAi* (gray) controls across baseline (0-36 hours), overnight mechanical sleep deprivation (36-48 hours), and recovery sleep (48-96 hours). Pink shading indicates time of sleep deprivation. Dotted lines = sleep deprivation, solid lines = undisturbed controls. (K) *;mex1-GAL4/+; UAS CG11037 RNAi/+* flies have less rebound sleep (ZT0-4) following 12 hours of mechanical sleep deprivation (pink bar on x-axis) compared to controls. (L) Enterocyte CG11037 knockdown flies have less rebound sleep during the full 12-hour light period following 12 hours of sleep deprivation. (M) Gut enterocyte *UAS CG11037 RNAi* flies have increased sleep latency after mechanical sleep deprivation compared to *GAL4* and *UAS* controls. C-E: Unpaired t-test, G-I, K-M: One way ANOVA with Tukey’s multiple comparisons. N > 4 biological replicates. The following are the numbers of flies (*n*) as plotted from left to right: **b-e;** *n =* 29, 32, **f-i;** *n =* 127, 132, 110, **j-m;** control: *n =* 16, 32, 16; sleep deprived: *n =* 16, 32, 16. C-E, G-I, K-M: Each dot represents one fly. B, F, J: Each line represents the mean sleep from 16 flies per genotype/condition. All columns are mean ± SEM; asterisks represent P-values *P≤0.05, **P≤0.01, ***P≤0.001, ****P≤0.0001, ns = not significant.

### *CG11037* is required in gut enterocytes for daily sleep regulation

We prioritized CG11037 for further study due to its robust and reproducible phenotype of sleep decrease. CG11037 encodes a trypsin-like serine endopeptidase; its knockdown specifically in the adult gut resulted in a loss of >220 minutes of sleep in both male and female flies (Figure 1B-C, Figure S1A-B). Analysis of sleep across the 24-hour cycle revealed that gut-specific CG11037 knockdown significantly reduced both daytime and nighttime sleep (Figure 1C, Figure S1B), with the night-time decrease manifesting largely as an early decline at the end of the night. These findings are consistent with a model in which CG11037-dependent proteolysis in the gut modulates peripheral signals that broadly influence sleep regulation.

Reduced sleep can be accounted for by a reduction in the total number of sleep bouts, shortened duration of individual sleep bouts, or a combination of both (21). To define the behavioral basis of sleep loss, we analyzed sleep architecture. CG11037 knockdown during adulthood increased latency to both day and night sleep (Figure 1D-E, Figure S1C-D, Table S3), suggesting impaired sleep initiation or reduced sleep drive. However, an increased number of shorter bouts at night is indicative of sleep fragmentation. During the day, bout number was reduced with no change in bout duration (Figure 1D-E, Figure 1C-D) and we found no change in waking activity (Table S3). Together, changes indicate a deficit in initiating sleep at the beginning of the night and subsequently in maintaining it.

The TiG2-S driver is expressed broadly in the gut (18). Single cell sequencing data indicate that CG11037 is expressed in gut enterocytes and enteroendocrine cells (22). To define the relevant cell type, we selectively knocked down *CG11037* in adult enterocytes using the temperature-sensitive TARGET system *(;mex1-GAL4, tub-GAL80^ts^*) (23). Enterocyte-specific knockdown during adulthood was sufficient to reduce sleep, recapitulating the phenotype observed with *TiGS-2* (Figure S1E-F). Consistent with this, constitutive knockdown using independent enterocyte drivers (*mex1-GAL4* and the split-GAL4 line *VT004958-AD; VT004958-DBD*) reduced both daytime and nighttime sleep, phenocopying *TiGS-2* mediated knockdown (Figure 1F-G, Figure S2A-B, 2E-F). In contrast, knockdown using two independent enteroendocrine cell drivers had no effect on sleep (Figure S2G-H). Together, these results demonstrate that *CG11037* is required in enterocytes of the midgut for baseline sleep regulation. Due to leakiness of the *TiGS-2* gene-switch line (Table S3), all subsequent gut-specific experiments were performed using the mex1-GAL4 driver.

Consistent with adult-only knockdown of *CG11037*, constitutive knockdown of *CG11037* in enterocytes produced a similar sleep architecture (Table S3), with an increase in daytime and nighttime latency (Figure 1H-I, Figure S2C-D). During the day, knockdown flies had shorter sleep bouts (Figure 1H-I, Figure S2C-D), whereas at night knockdown flies had shorter bouts but many more of them (Figure 1H-I, Figure S2C-D). Thus, like adult only knockdown, *CG11037* constitutive knockdown flies have delayed onset and fragmented sleep at night.

Independent mechanisms regulate sleep under undisturbed conditions, and homeostatic sleep rebound following deprivation (24,25). To determine if sleep homeostasis is intact in *CG11037* deficient flies, we sleep deprived flies via random mechanical shaking for 12-hours throughout the night and measured sleep the following day. This sleep deprivation protocol was sufficient to sleep deprive all flies (Figure 1J, pink bar). *CG11037* knockdown flies had significantly less rebound sleep following sleep deprivation (Figure 1K-L) and increased latency to sleep (Figure 1M). Although the reduced rebound could derive from reduced loss of sleep given the lower baseline in these flies, the increased latency supports the idea that *CG11037* gut knockdown flies do not accumulate as much sleep need as controls when they are deprived.

To exclude off-target RNAi effects, we validated the behavioral effects of *CG11037* knockdown using two independent RNAi lines and *CG11037* loss-of-function mutants as a result of a transposable element insertion into the promoter of the gene (26). All lines recapitulated the reduced sleep phenotype relative to controls (Figure 3A-C). Furthermore, qPCR analysis of dissected guts confirmed that each RNAi and mutant line significantly reduced *CG11037* transcript levels (Figure 3D-F), supporting the specificity of the observed phenotype.

To test sufficiency of gut expression of *CG11037* for sleep, we restored *CG11037* expression in the enterocytes of the midgut in *CG11037* mutants. Gut-specific (*;mex1-GAL4/+; CG11037 mutant/UAS CG11037*) restored total sleep, bout number, bout length, and sleep latency to control levels in both sexes (Figure S3G, Table S3). In contrast, overexpression of *CG11037* in the gut of wild-type flies either throughout life or only during adulthood (Figure S3H-J) did not increase sleep, indicating that *CG11037* is necessary for sleep, but not sufficient to promote sleep beyond baseline levels.

Given that serine endopeptidases can shape microbial composition (27,28) and microbial metabolite production is under circadian regulation (29,30) and may influence sleep behavior (31,32), we sought to determine if *CG11037* acts through the microbiome to modulate sleep. We reared *CG11037* gut knockdown flies and control flies under control or microbiome-depleted conditions and measured sleep. To confirm that microbiome depleted flies were in fact microbiome free, siblings were tested for growth of several bacterial species using different media growth plates (methods and data not shown). We found that sleep loss persisted in microbiome depleted *CG11037* RNAi flies (Figure S4). These findings suggest that *CG11037* regulates sleep through host-intrinsic mechanisms rather than through alterations in the microbiota.

Although *CG11037* is not expressed in brain according to publicly available single-nucleus RNA sequencing data (33), we confirmed that targeting CG11037 RNAi pan neuronally (*nSyb-GAL4*) does not affect sleep (Figure S5).

### *CG11037* knockdown in the gut affects the response to sickness

Serine endopeptidases in the gut are key regulators of epithelial homeostasis (34,35) and innate immune signaling (36,37), functioning upstream of NF-kB-dependent pathways (38). Sleep in Drosophila is sensitive to immune state, in particular following bacterial infection or sterile injury (39). We hypothesized that loss of gut protease *CG11037* might alter sickness sleep by modulating basal immune tone or host-intrinsic infection responses.

To test this, we subjected gut-specific *CG11037* knockdown flies to bacterial infection using *S. marcescens*, a gram negative bacteria species, at ZT18, an infection time that produces an increase in sleep the following morning(39,40). While control flies exhibited the expected increase in sleep at ZT0-4, *CG11037* knockdown flies did not show additional sleep induction (Figure 2A-B). Despite the loss of sickness sleep, *CG11037* knockdown flies survived *S. marcescens* infection at rates comparable to control animals (Figure 2C), demonstrating that host defenses are not compromised. Indeed, bacterial growth was more effectively restricted 24 hours post-infection compared to controls (Figure 2D), indicating that knockdown animals are better at clearing infection. We also performed sterile injury with PBS at ZT18, and as with bacterial infection, did not observe sickness sleep in *CG11037* knockdown flies the following day (Figure 2E-F). Given that sickness sleep is normally restorative and *CG11037* flies show robust survival, these data suggest that loss of *CG11037* reduces the need for sleep to recover from sickness.

**Figure 2:**
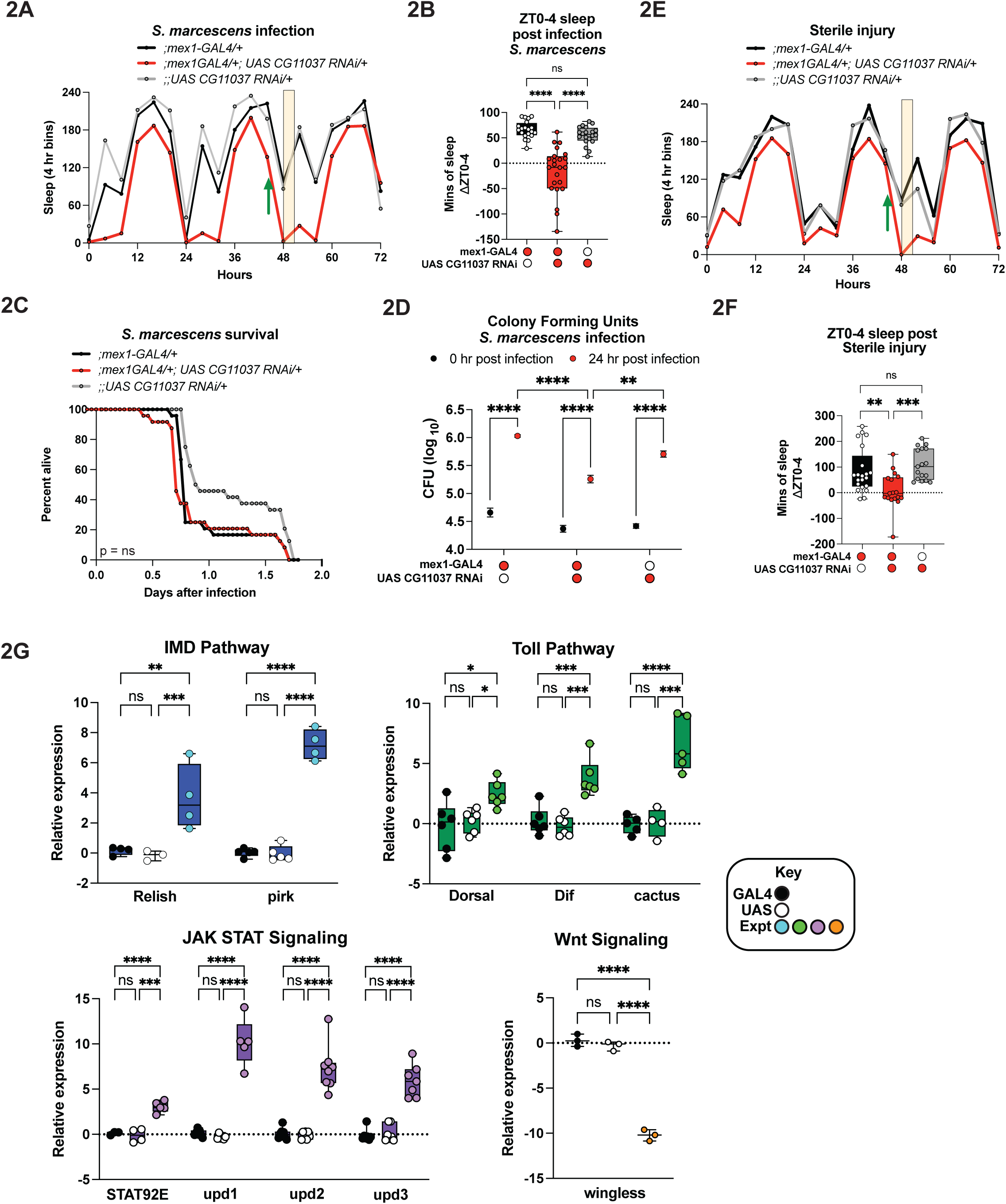
Loss of *CG11037* uncouples sickness sleep from survival. (A) Representative sleep trace from infection with *S. marcescens* at ZT 18 (green arrow). Mean sleep is plotted for 3 consecutive days in 4 hr increments. Shaded area indicates the increase in sleep from ZT0-4 after infection. (B) Compared to the *GAL4* and *UAS* controls, whole life gut knockdown of *CG11037* eliminates sickness sleep typically seen ZT0-4 following infection. (C) There is no difference in survival following *S. marcescens* infection. (D) Following 24 hours of *S. marcescens* infection bacterial load is lower in *;mex1-GAL4/+; UAS CG11037 RNAi/+* flies compared to *GAL4* and *UAS* controls. (E) Representative sleep trace from sterile injury with PBS at ZT 18 (green arrow). Mean sleep is plotted for 3 consecutive days in 4-hour increments. Shaded area indicates the increase in sleep from ZT0-4 after injury. (F) *mex1-GAL4/+; UAS CG11037 RNAi/+* flies do not engage in sickness sleep following ZT18 sterile injury compared to controls. (G) Relative RNA expression level measured by qPCR in isolated dissected guts of *;mex1-GAL4/+; UAS CG11037 RNAi/+* (colored circles) compared to *;mex1-GAL4/+* controls (white circles) or *;;UAS CG11037 RNAi/+* controls (black circles). B, F: One way ANOVA with Tukey’s multiple comparisons, C: Kaplan-Meier with Log-rank test, D: Two-way ANOVA with Tukey’s multiple comparisons, G: One-way ANOVA with Turkey’s multiple comparisons. N > 3 biological replicates. The following are the numbers of flies (*n*) as plotted from left to right: **a-c**: *n* = 20, 23, 20; **d:** *n =* 60 crushed flies per condition; **e-f:** *n =* 21, 17, 17. B and F: Each dot represents one fly. A, C, E: Each line represents the mean sleep/survival of the number of flies indicated above. G: each dot represents one biological replicate which is the average of three technical replicates using at minimum 35 dissected guts. All columns are mean ± SEM; asterisks represent P-values *P≤0.05, **P≤0.01, ***P≤0.001, ****P≤0.0001, ns = not significant.

As CG11037 knockdown flies survive infection comparable to control flies, despite not having sickness sleep, we asked whether immune signaling pathways were altered under basal conditions. To address this, we assayed expression of components of the Drosophila Immune Deficiency (Imd), Toll, JAK-STAT, and c-Jun N-terminal kinase (JNK) pathways using qPCR. *CG11037* knockdown flies displayed elevated expression of *Relish* and the IMD negative regulator *Pirk* (41) relative to both GAL4 and UAS controls (Figure 2G), suggesting enhanced basal activity of the IMD pathway. We next examined the Toll signaling pathway, which responds primarily to Gram-positive bacteria (42) and fungi (43). Like the IMD pathway, *CG11037* knockdown flies showed increased transcript levels of *Dorsal*, *Dif*, and the pathway inhibitor *Cactus* (Figure 2G), indicating that multiple NF-κB-dependent immune pathways are transcriptionally elevated. We next assessed the JAK-STAT signaling pathway, which is commonly activated during viral infection (44) and epithelial stress (45). CG11037 gut knockdown flies exhibited increased expression of all three pathway ligands *upd1*, *upd2*, and *upd3* as well as the transcription factor *Stat92E* (Figure 2G), suggesting increased cytokine signaling in the gut. Finally, we examined components of the JNK signaling pathway and found reduced expression of *wingless* in *CG11037* knockdown flies compared to controls (Figure 2G). Together, these results indicate that loss of *CG11037* leads to broad changes in immune and stress signaling pathways at baseline.

### *CG11037* knockdown alters the proteome

To identify potential targets of CG11037 that may contribute to sleep changes, we performed LC-MS/MS on bodies (with heads, legs, and wings removed) of 4–6-day old female *GAL4/+* and *UAS/+* controls and *;mex1-GAL4/+; UAS CG11037 RNAi/+* and independently on extracted hemolymph. Because CG11037 is a predicted serine endopeptidase, samples were analyzed without digestion by Trypsin to preserve endogenous proteolytic fragments. Under these conditions, detected peptides are expected to reflect naturally occurring proteolytic products. We reasoned that proteins changed in abundance in *CG11037* knockdown animals may represent candidate targets.

Prior to collection of tissue, siblings were phenotypically verified for sleep decreases (data not shown). Comparative analysis of bodies identified 47 proteins significantly altered between controls and *CG11037* gut knockdown flies (FDR < 0.25, Figure 3A). Of these, 25 were increased and 22 were decreased in knockdown animals (Figure 3A). On the other hand, comparative analysis of hemolymph samples identified 22 proteins significantly altered between control and *CG11037* gut knockdown flies (FDR <0.25, Figure 3B). Of these 19 were increased and 3 were decreased in knockdown flies (Figure 3B).

**Figure 3:**
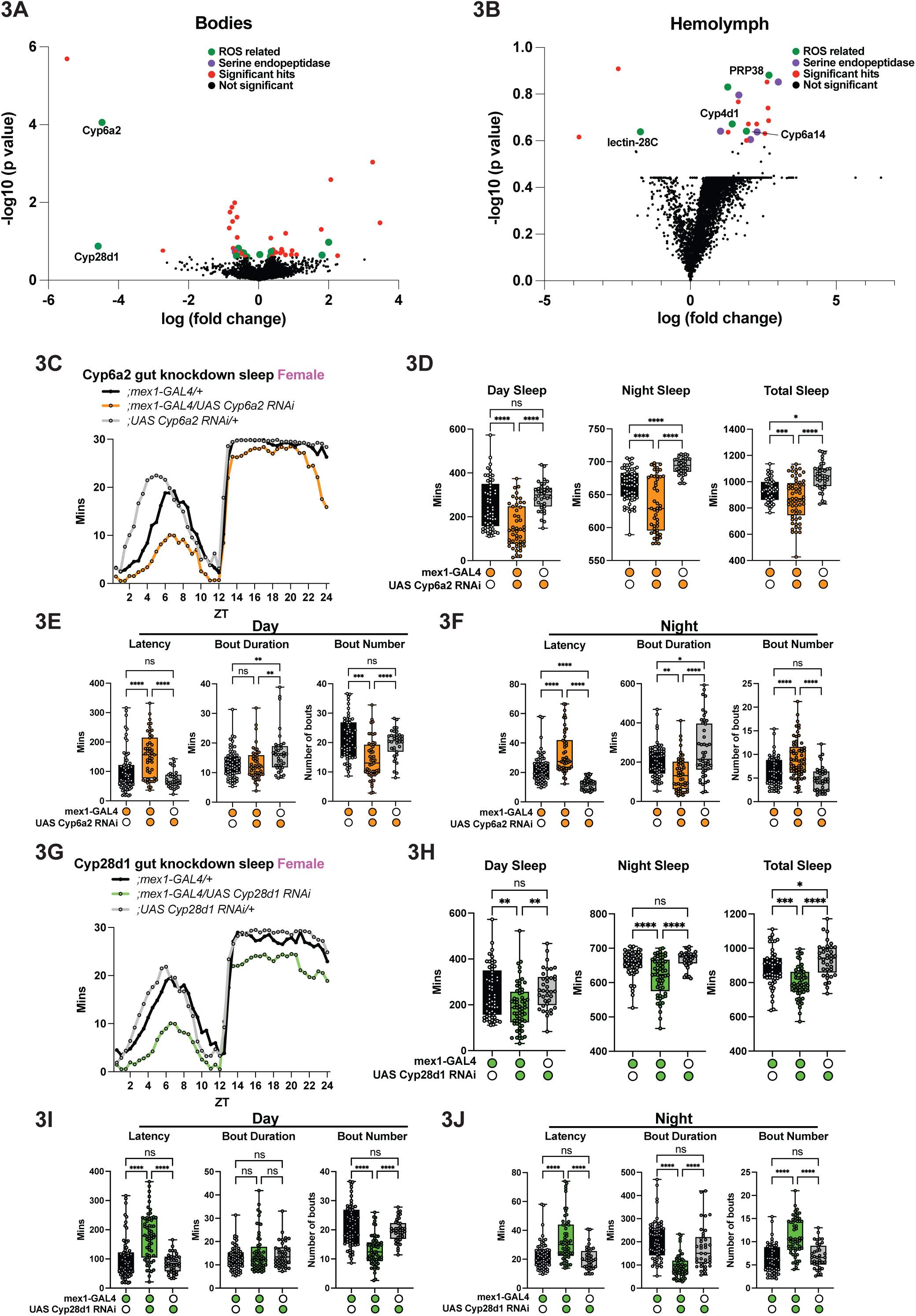
Loss of *CG11037* alters expression of ROS-regulating proteins. (A-B) Proteomic data comparing bodies (A) or hemolymph (B) of gut enterocyte specific knockdown flies *(;mex1-GAL4/+; UAS CG11037 RNAi/+*) to *UAS* and *GAL4* control bodies. Differentially expressed proteins are colored in green (ROS related), purple (serine endopeptidase), red (other significant hits), while not significant proteins are labeled in black. 47 differentially expressed proteins were identified in body samples, and 19 identified in hemolymph samples. Normalized log expression counts are shown for all proteins with a p-value <0.25 across all fold changes. (C) Representative sleep trace of *;mex1-GAL4/UAS Cyp6a2 RNAi* (orange), and *UAS* (gray) and *GAL4* (black) controls. (D) Whole life enterocyte knockdown of *Cyp6a2* decreases day (left), night (middle), and overall sleep (right). (E) Gut knockdown of *Cyp6a2* increases daytime latency, does not affect bout duration, but decreases the number of daytime sleep bouts. (F) At night *;mex1-GAL4/UAS Cyp6a2 RNAi* increases sleep latency, reduces bout duration, while increasing the number of night sleep bouts. (G-H) Gut knockdown of *Cyp28d1* (green) reduces day and night sleep, resulting in an overall short sleep phenotype. (I) Gut enterocyte knockdown of *Cyp28d1* increases daytime latency, without affecting bout length. However, *Cyp28d1* knockdown decreases overall day time bout number. (J) At night *;mex1-GAL4/UAS Cyp28d1 RNAi* increases sleep latency, decreases bout length, and increases overall bout number. D-F, H-J: One way ANOVA with Tukey’s multiple comparisons. For panels A and B, 6 biological replicates, All other panels N > 4 biological replicates. The following are the numbers of flies (*n*) as plotted from left to right: **c-f:** *n* = 63, 48, 47; **e-h:** *n =* 63, 48, 47. D-F, H-J: Each dot represents one fly. C and G: Each line represents the mean sleep from 16 flies per genotype. All columns are mean ± SEM; asterisks represent P-values *P≤0.05, **P≤0.01, ***P≤0.001, ****P≤0.0001, ns = not significant.

Functional categorization of the differentially expressed proteins revealed enrichment for factors associated with oxidative stress regulation. Of the 47 proteins altered in body samples, 11 are annotated as regulators of redox homeostasis (5/11 down regulated, 6/11 upregulated), representing nearly one quarter of the identified proteome. Oxidative stress related proteins included the UPD-glycosyltransferase family 37 members D1 (Ugt37D1), a detoxification enzyme that participates in xenobiotic metabolism and oxidative stress responses(46), the glutathione peroxidase homolog Gtpx, a thioredoxin-dependent peroxidase that protects cells from oxidative damage (47), and metabolic enzymes such as fatty acid synthase 3 (FASN3), which has been linked to lipid metabolic pathways that influence cellular redox balance (48). In addition, sanpodo (spdo), previously reported to be regulated in response to arsenite-induced stress (49), was also identified in this dataset. Among the most strongly downregulated proteins in this category were Cytochrome P450 6a2 (Cyp6a2) and Cytochrome P450 28d1 (Cyp28d1), two oxidative stress-associated enzymes that we selected for further analysis as described below. Consistent with the presence of oxidative stress-associated factors in the dataset, 5/19 differentially expressed proteins in hemolymph samples were linked to oxidative stress responses. Interestingly, while Cytochrome P450 proteins are decreased in abundance in body samples, we find that different Cytochrome P450s are increased in expression in the hemolymph (Figure 3B). In addition, 5/19 differentially regulated hemolymph proteins were serine endopeptidases.

### Gut-specific knockdown of *Cytochrome P450s* phenocopies *CG11037* knockdown

Cyp6a2 and Cyp28d1 were among the Cytochrome P450 proteins whose abundance in bodies was significantly reduced following gut-specific knockdown of *CG11037*. This is mechanistically compelling for two reasons. First, Cytochrome P450 expression is tightly regulated by immune and stress signaling pathways (50,51). Second, proteomic analysis revealed alterations in multiple proteins involved in oxidative stress regulation, suggesting that CG11037 influences pathways that control redox homeostasis. Cyp6a2 and Cyp28d1 participate in detoxification and oxidative stress responses (52–54) in *Drosophila* and consistent with a role for them here, they are expressed in enterocytes (22). Together, these observations provided a rationale to investigate Cyp6a2 and Cyp28d1 as potential mediators of the sleep phenotype observed following *CG11037* knockdown.

We asked if enterocyte/mid gut-specific knockdown of *Cyp6a2* or *Cyp28d1* would phenocopy *CG11037* knockdown and found that adult-only gut knockdown of *Cyp6a2* and *Cyp28d1* decreased both male and female sleep (Figure S6A-B). As with *CG11037*, constitutive knockdown of *Cyp6a2* or *Cyp28d1* in enterocytes also decreased sleep (Figure 3C-D, 3G-H, Figure S6C-D, 6G-H). Like the sleep trace of enterocyte *CG11037* knockdown, *Cyp6a2* and *Cyp28d1* gut knockdown flies show a decline in sleep starting at ZT19 although *Cyp28d1* females showed generally reduced sleep throughout the night (Figure 3G). Gut knockdown of *Cyp6a2* or *Cyp28d1* also changed sleep architecture, with increased sleep latency in the day and night for all except *Cyp6a2* knockdown males during the day (Figure 3E-F, 3I-J, Figure S6E-F, 6I-J, Table S3). Similar to *CG11037* knockdown, *Cyp6a2/Cyp28d1* knockdown flies had fewer daytime sleep bouts with no change in duration (Figure 3E-F, 3I-J, Figure S6E-F, 6I-J, Data S3), but increased night-time sleep bouts that were shorter in length (Figure 3E-F, 3I-J, Figure S6E-F, 6I-J, Table S3). Changes in sleep were not accompanied by changes in waking activity levels (Table S3).

### Knockdown of *CG11037*, *Cyp6a2*, or *Cyp28d1* increases ROS levels in the gut but not the brain

Proteomic analysis following serine endopeptidase knockdown revealed coordinated reductions in multiple Cytochrome P450 enzymes and other known redox regulators. Cytochrome P450s are major regulators of cellular redox balance. Cytochrome P450s, including both Cyp6a2 and Cyp28d1, contribute to xenobiotic detoxification and metabolic buffering (54–56). The convergence of these redox-active systems strongly predicts altered reactive oxygen species (ROS) levels. Moreover, oxidative stress has been implicated in sleep regulation (57,58).

We hypothesized that gut knockdown of *CG11037* or Cytochrome P450s would increase ROS activity in the gut. To test this, Dihydroethidium (DHE) was used as a specific dye to measure ROS levels in freshly dissected female guts. Total DHE fluorescence was higher in the gut of *;mex1-GAL4/+; UAS CG11037 RNAi/+* flies compared to controls (Figure 4A-B). DHE fluorescence was highest in the midgut and slowly decreased toward the anterior and posterior ends of the gut. Similarly, we found that RNAi of *Cyp6a2* or *Cyp28d1* increased DHE fluorescence in the gut (Figure 4C-F). Elevated reactive oxygen species can be genotoxic, so we checked for DNA damage in the guts of knockdown flies but found no difference in damage levels (Figure S7A). Together, this suggests that *CG11037*, *Cyp6a2*, and *Cyp28d1* normally function to maintain low levels of ROS in the intestine.

**Figure 4:**
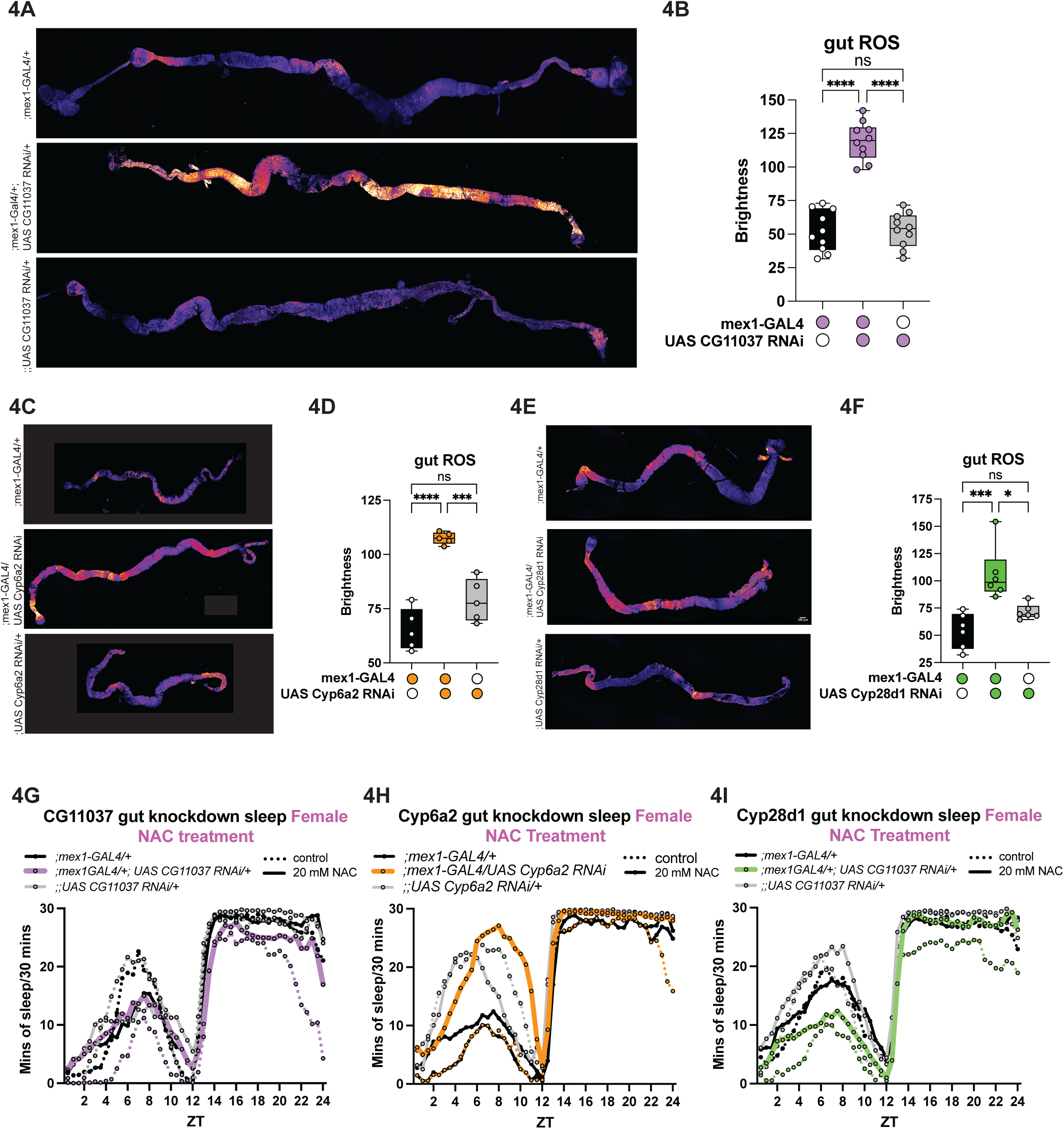
Elevated gut ROS drive sleep loss following *CG11037* and *Cytochrome P450* knockdown. (A) Representative images of Reactive Oxygen Species (ROS) accumulation in the gut of enterocyte *CG11037* knockdown flies (middle), *UAS* and *GAL4* controls. (B) ROS accumulation is higher in gut *CG11037* knockdown flies (purple) compared to controls. (C) Representative images of baseline ROS accumulation in dissected guts of gut *Cyp6a2* knockdown, *UAS* and *GAL4* control flies. (D) Quantification of DHE staining suggests that gut knockdown of *Cyp6a2* (orange) increases baseline levels of ROS compared to controls. (E) Representative images of DHE stained *;mex1-GAL4/UAS Cyp28d1 RNAi*, *UAS* and *GAL4* control dissected guts. (F) Quantification of DHE staining shows that gut knockdown of *Cyp28d1* increases baseline levels of ROS. (G) NAC Antioxidant treatment (solid lines) is sufficient to restore sleep in *CG11037* gut knockdown flies (purple lines). (H) Short sleep of enterocyte *Cyp6a2* knockdown (orange lines) is ameliorated by antioxidant treatment via NAC. (I) Antioxidant treatment increases sleep in *Cyp28d1* gut knockdown flies (green lines). B, D, F: one way ANOVA with Tukey’s multiple comparisons. N > 3 biological replicates. Each imaging experiment used at minimum 5 dissected guts, and sleep experiments used at minimum 16 flies per genotype per treatment. B, D, F: Each dot represents one fly. G-I: Each line represents the mean sleep from 16 flies per genotype per condition. All columns are mean ± SEM; asterisks represent P-values *P≤0.05, ***P≤0.001, ****P≤0.0001, ns = not significant.

Because these observations suggested a role for CG11037 in regulating oxidative stress, we next asked whether *CG11037* itself is responsive to oxidative stress. Wild-type flies were exposed to 3% hydrogen peroxide, and *CG11037* expression was measured. We found that *CG11037* mRNA levels increased after the treatment (Figure S7B), consistent with a potential role in oxidative stress regulation.

We next exposed *;mex1-GAL4/+; UAS CG11037 RNAi/+* flies and control flies to 3% hydrogen peroxide for 24 hours and subsequently measured ROS levels. We found that control flies increased DHE fluorescence following oxidative stress challenge (Figure S7C). However, *CG11037* knockdown flies did not increase DHE fluorescence further. Next, we treated hydrogen peroxide treated flies with 20 mM N-acetyl lysine (NAC) as an antioxidant to reduce oxidative stress. We found that 24 hours of 20 mM NAC treatment was sufficient to reduce ROS stress in all groups (Figure S7C).

*CG11037* exerts its sleep function in the periphery, but sleep is ultimately executed through brain circuits (8). We hypothesized that *CG11037* may exert its function by altering ROS levels in the brain. We used DHE staining on freshly dissected female brains to determine if ROS levels were altered. Unlike gut ROS, we found no differences in brain ROS in *CG11037* gut knockdown flies (Figure S7D).

### Gut ROS underlies reduced sleep in *CG11037* and *Cyp450* knockdown flies

It was previously reported that short-sleeping animals are hypersensitive to oxidative stress (59–61). We compared the survival of *CG11037 RNAi* flies relative to controls when subjected to two different treatments that induce oxidative stress by increasing ROS levels. We exposed flies to paraquat and immediately began measuring survival. Although there was variability in the paraquat response even among the controls, *CG11037 RNAi* flies were not significantly different from control flies (Figure S7E). In other experiments, *CG11037* knockdown flies and controls were fed 4% hydrogen peroxide, an oxidant that produces highly reactive hydroxyl radicals. Unlike with paraquat treatment, *CG11037* knockdown flies were more sensitive than controls to hydrogen peroxide (Figure S7F). Thus, *CG11037* knockdown flies are susceptible to some kinds of oxidative stress.

*CG11037* gut knockdown flies are short sleepers with elevated levels of intestinal ROS. To determine whether ROS contributes to short sleep in these flies, we reduced whole body ROS levels by exposing flies to 20 mM NAC for 48 hours prior to measuring sleep and continued treatment throughout the sleep analysis period. DHE fluorescence had already indicated that feeding 20 mM NAC was sufficient to bring *;mex1-GAL4/+; UAS CG11037 RNAi/+* gut ROS levels down to those of wild-type flies (Figure S7C) so it provided an effective method to evaluate the effect of ROS on behavior. We found that 20 mM NAC treatment was sufficient to increase *CG11037* deficient sleep back to control levels during the day and night (Figure 4G, Figure S8A).

Our data thus far suggest that loss of *CG11037* in the gut reduces Cytochrome P450 levels, thereby increasing peripheral ROS levels, which leads to reduced sleep. If this is the case, *Cyp6a2* and *Cyp28d1* RNAi short sleep should also be ROS dependent. We pre-treated *Cyp6a2* and *Cyp28d1* RNAi and control flies with 20 mM NAC for 48 hours and then assayed sleep while flies remained on 20 mM NAC. As with gut *CG11037* knockdown, sleep loss in *Cyp6a2* or *Cyp28d1* knockdowns was rescued back to wild-type levels with antioxidant treatment (Figure 4H-I, Figure 8B-C). Together these results demonstrate that both *CG11037* and Cytochrome P450s modulate sleep amount via ROS dependent pathways.

### *CG11037* and *Cytochrome P450* knockdown flies are long-lived

*CG11037* knockdown flies exhibit reduced sleep and yet do not alter susceptibility to infection despite improved resistance leading us to ask how loss of this gut protease affects lifespan. Strikingly, gut-specific *CG11037* knockdown extended both male and female lifespan compared to control flies (Figure 5A, Figure S9A). To determine whether this longevity was linked to redox balance, we treated animals with 20 mM NAC, which suppressed the lifespan extension (Figure 5B, Figure S9B). Additionally, we assayed lifespan following knockdown of *Cyp6a2* or Cyp28d1 in the gut and found that these animals are also long-lived (Figure 6C, 6E, Figure S9C, 9E). Similarly, antioxidant treatment suppressed this lifespan extension (Figure 6D, 6F, Figure S9D, 9F). These findings indicate that knockdown of CG11037 or of its putative downstream targets, *Cytochrome P450 6a2* and *28d1*, promotes ROS-dependent lifespan extension.

**Figure 5:**
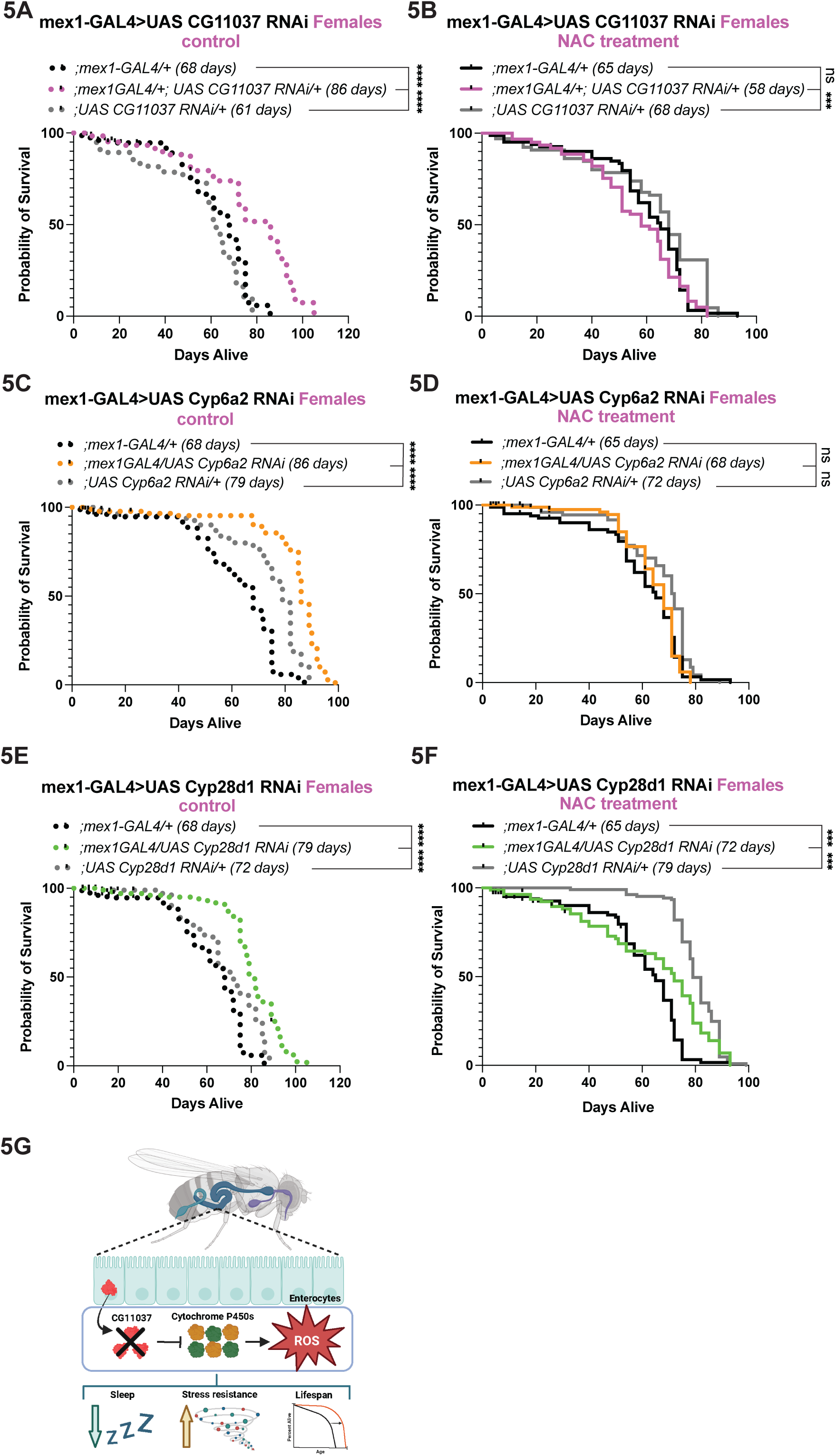
Lifespan extension by *CG11037* and *Cytochrome P450* is dependent on ROS. (A) Enterocyte knockdown of *CG11037* extends lifespan compared to *GAL4* and *UAS* controls. (B) Treatment with antioxidants (NAC) throughout life suppresses lifespan extension in *;mex1-GAL4/+; UAS CG11037 RNAi/+* flies. (C) Constitutive gut knockdown of *Cyp6a2* extends lifespan compared to *UAS* and *GAL4* controls. (D) Lifespan extension by gut *Cyp6a2* knockdown is dependent on ROS accumulation as antioxidant treatment suppresses lifespan extension. (E) Enterocyte knockdown of *Cyp28d1* extends lifespan compared to *GAL4* and *UAS* controls. (F) Lifespan extension in *;mex1-GAL4/ UAS Cyp28d1 RNAi* is ROS dependent as it is suppressed by antioxidant treatment. (G) Model: Gut/enterocyte knockdown of CG11037 (red) decreases expression of Cytochrome P450s Cyp6a2 (green) and Cyp28d1 (orange) resulting in excess ROS accumulation. Excess ROS leads to a decrease in sleep, increase in stress resistance, and an extended lifespan. Median survival is displayed next to genotypes. A-F: Kaplan Meier with Log-rank test. N = 3 biological replications. Each lifespan begins with minimum 80 flies. Asterisks represent P-values ***P≤0.001, ****P≤0.0001, ns = not significant.

## Discussion

Sleep has traditionally been viewed as a brain-driven process, yet growing evidence indicates that peripheral physiology can influence sleep behavior (62,63). Using a targeted screen of secreted peptides expressed in peripheral tissues, we discovered that the trypsin-like serine endopeptidase *CG11037* functions in midgut enterocytes to maintain normal sleep duration and architecture. Gut loss of *CG11037* reduces normative sleep and sickness sleep responses, elevates gut reactive oxygen species (ROS), and alters the abundance of Cytochrome P450 enzymes and other stress response proteins. Importantly, despite their reduced sleep, flies lacking *CG11037* in the gut have robust defense responses and increased lifespan, and sleep and lifespan are attributable to the higher gut ROS (Figure 5G). We propose that the stress adaptation of these animals’ accounts for their reduced sleep and extended longevity and suggest a mechanism by which moderate elevations in stress signals including oxidative stress can be beneficial.

### A gut protease pathway controls sleep amount and architecture

Although neural circuits that control sleep have been extensively characterized (8,64), the mechanisms through which peripheral tissues regulate sleep remain poorly understood. Most prior studies have examined candidate hormones or immune factors, often under conditions of acute stress. By contrast, our screen of the peripheral secretome represents a relatively unbiased effort to identify molecules through which peripheral tissues influence normal daily sleep. This approach identified a previously unrecognized class of molecules, serine endopeptidases, as regulators of daily sleep. The enrichment of trypsin/chymotrypsin-like proteases among sleep-regulating hits suggests that extracellular proteolytic processing may broadly shape systemic signals that influence sleep.

We demonstrate that *CG11037* is specifically required in enterocytes to maintain consolidated sleep. Loss of *CG11037* produces a distinctive sleep phenotype characterized by a drop off in sleep in the second half of the night. Together with the increased latency to sleep at night and following deprivation, these data suggest that *CG11037* knockdown animals may be short sleepers with low sleep need.

The expression of *CG11037* in enterocytes is notable because these cells represent the primary metabolic interface between the organism and its environment. Enterocytes coordinate nutrient absorption, detoxification, immune responses, and metabolic signaling, processes that can influence systemic physiology (65–68). Our findings therefore suggest that metabolic and immune processes occurring within intestinal epithelial cells can influence sleep.

### Cytochrome P450 enzymes link gut protease activity to redox metabolism

Proteomic analysis revealed that loss of *CG11037* alters expression of Cytochrome P450 enzymes, including Cyp6a2 and Cyp28d1. Cytochrome P450s are central components of metabolic detoxification pathways and play important roles in maintaining cellular redox (52,54,69). Gut-specific knockdown of either Cyp6a2 or Cyp28d1 phenocopies the short-sleep and sleep architecture observed in CG11037-deficient flies, supporting the idea that these enzymes mediate effects of CG11037.

One possibility is that CG11037 directly regulates P450 activity through proteolytic processing. In silico analyses identified specific trypsin cleavage motifs compared to other enzymatic cleavage motifs within several cytochrome P450 enzymes, including both Cyp6a2 and Cyp28d1 (Expasy). Alternatively, CG11037 may regulate upstream signaling pathways that control P450 levels. Cytochrome P450 expression is highly responsive to immune activation, oxidative stress, and xenobiotic signaling, all of which could be influenced by intestinal protease activity (52,54,69). Although the precise molecular relationship between CG11037 and P450 regulation remains to be determined, these findings suggest that intestinal proteolytic signaling influences metabolic pathways that maintain redox homeostasis.

### Peripheral ROS as a regulator of sleep

Our data identify reactive oxygen species (ROS) as a key mechanistic link between gut/peripheral physiology and sleep behavior. Knockdown of *CG11037* or downstream P450 enzymes elevates ROS specifically in the gut without altering ROS levels in the brain. Importantly, reducing systemic oxidative stress using an antioxidant restores sleep amount, demonstrating that elevated ROS is required for the short-sleep phenotype.

Oxidative stress has previously been linked to sleep regulation in Drosophila and mammals. Sleep deprivation increases markers of oxidative damage(57,70,71), and antioxidant pathways can influence host physiology in the face of sleep loss(72,73). Although high levels of intestinal oxidative stress resulting from extreme sleep loss (90% sleep loss) was shown to account for mortality, this oxidative damage was not implicated in changes in sleep (70). Our findings suggest that moderate increases in ROS can cause moderate loss of sleep (∼25%), perhaps through circulating metabolites, immune regulators, or neuroendocrine factors released from the gut epithelium. Alternatively, ROS-dependent signaling could alter vagal-like sensory pathways that convey a visceral physiological state to the brain.

### Short sleep with extended lifespan

Sleep is recognized as crucial for health and longevity, with short-sleeping animals typically showing a reduced lifespan. There are examples of naturally short-sleeping humans who are not compromised by sleep loss, but the underlying mechanisms are not known. Strikingly, *CG11037* knockdown flies exhibit extended lifespan despite reduced sleep and increased oxidative stress. This may relate to findings that low levels of oxidative stress can promote internal homeostasis and stress resistance (74–78), even though extreme levels of ROS lead to mortality potentially through excess DNA damage and apoptosis (70). The phenotype reported here, moderate levels of gut ROS accompanied by moderate sleep loss (∼25% reduction). with no DNA damage, resembles that of several stress-adapted longevity models such as the naked mole rat (24), Bowhead whales (79,80), insulin signaling mutants across species (81,82), and mitochondrial dysfunction models including *clk-1* (83,84) and *isp-1* (85) mutant *C.* elegans and *Surf1* knockout mice in which mild chronic stress promotes lifespan extension through hormetic mechanisms (86,87). Here we show that *CG11037* knockdown animals, at baseline, have higher levels of some of these pathways including IMD, Toll, and JAK-STAT signaling. Similarly, in long-lived models, including the naked mole rat and mitochondrial dysfunction models, low levels of ROS activate protective transcriptional programs that enhance stress resistance and longevity. Consistent with this possibility, the lifespan extension observed in *CG11037*-deficient flies is ROS dependent, as antioxidant treatment abolishes the longevity phenotype.

An increased lifespan in a short-sleep fly mutant is quite surprising, as typically the sleep reducing mutants dramatically reduce lifespan (88–93). CG11037 flies even show a lack of sleep in response to immune challenges, despite which their survival/recovery is equivalent to that of wild type. We propose that these flies need less sleep, in other words that the mechanisms that prolong lifespan also allow them to function optimally with lower levels of sleep.

It is paradoxical that *CG11037* would have evolved given that its loss in the gut reduces sleep need and extends lifespan. There could be other untested detriments resulting from its loss such as sensitivity to some stressors, altered metabolism and energy expenditure. Indeed, we find that *CG11037* knockdown animals are sensitive to some inducers of oxidative stress, which could exist in the fruit flies’ natural environment as byproducts of fermenting fruit, including ethanol, acetaldehyde, organic acids and other microbial metabolites. In addition, some immune pathways transcriptionally upregulated upon *CG11037* knockdown, including *JAK-STAT* signaling in *CG11037* knockdown flies, could have negative effects on physiology. We find too that knockdown flies have decreased mRNA levels of wingless, the key ligand for stress resistance through Wnt signaling. Additionally, *CG11037* regulates Cytochrome P450 enzymes that play critical roles in detoxification and stress responses. In fluctuating environments where flies encounter dietary toxins, microbes, and other stressors, maintaining robust detoxification pathways may be essential for survival and reproduction. Thus, beneficial effects of *CG11037* loss may be specific to benign conditions.

Together our findings support a model in which the enterocyte protease CG11037 regulates intestinal redox balance through cytochrome P450 enzymes to maintain optimal levels of ROS, and thereby normal sleep and lifespan. Loss of *CG11037* disrupts this pathway, leading to elevated gut ROS. Increased intestinal oxidative stress then alters systemic physiological signaling, resulting in reduced sleep, altered immune-coupling, and ROS-dependent lifespan extension. In this model, the gut functions not only as a metabolic organ but also as an active regulator of behavioral state. Together, these findings highlight the importance of peripheral physiological states in shaping sleep, immunity, and aging.

## Acknowledgements

The authors would like to thank Sehgal lab members for their discussions and advice.

## Author Contributions

RSM and AS conceived the study. RSM, RX, and EA developed and validated methodology as needed. RSM and RX performed most experiments with assistance from EA, JW, TGB, and JS. All brains were dissected by EA. RSM analyzed most experiments with assistance from RX. LS, HF and EN performed and analyzed mass spectrometry data. RSM wrote and designed all graphics for the manuscript. RX and AS revised the manuscript. RSM and AS secured funding for this work. All authors contributed to intellectually to the manuscript.

## Competing Interesting

The authors declare no competing interests.

---

**Extended Data Figure 1:**
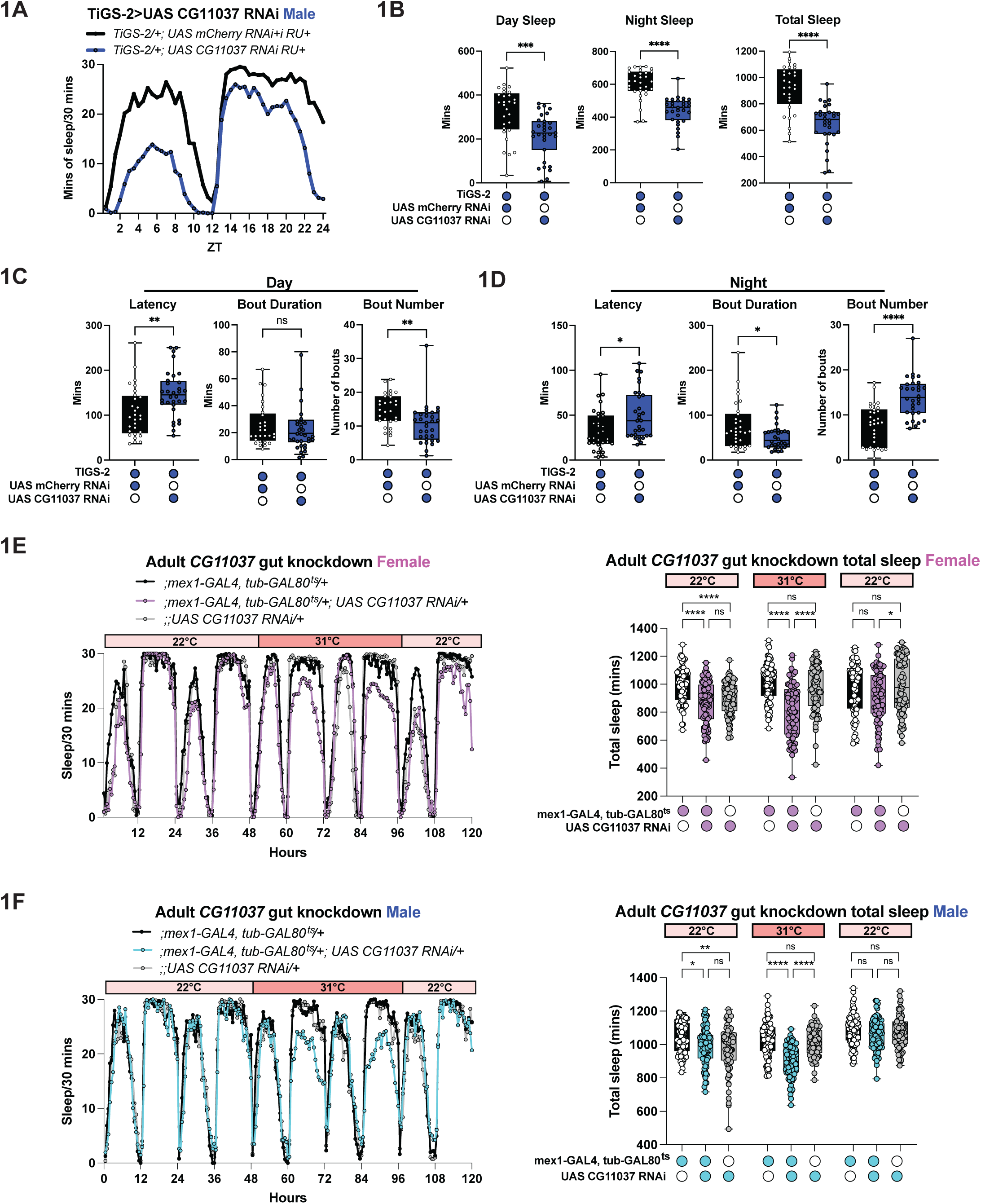
*CG11037* is required during adulthood in enterocytes for daily sleep. (A) Representative sleep trace of male *;TiGS-2/+; UAS CG11037 RNAi/+* (blue) compared to *;TIGS-2/+; UAS mCherry RNAi/+* (black) controls. (B) Adult only knockdown of *CG11037* using the *TiGS-2* gene switch driver decreases daytime (left) and nighttime (middle) resulting in overall less sleep (right). (C) Adult gut knockdown of *CG11037* increases daytime sleep latency (left), does not affect daytime bout duration (middle), but leads to fewer overall day sleep bouts (right). (D) Adult gut knockdown of *CG11037* increases nighttime sleep latency (left), decreases nighttime bout duration (middle), and increases nighttime bouts (right). (E-F) Representative sleep trace (left) of females (E) and males (F) in which CG11037 is knocked down only during adulthood using the TARGET system *(;mex1-GAL4, tub-GAL80^ts^/+; UAS CG11037 RNAi/+*) and controls. Quantified on the right; Sleep was recorded for 48 hours at 22°C, the temperature was then raised to 31°C to inactivate the GAL80^ts^ to achieve *CG11037* knockdown for two days, and finally temperature was lowered back to 22°C. B-D: Unpaired t-test, E-F: one way ANOVA with Tukey’s multiple comparisons test. N > 4 biological replicates. The following are the numbers of flies (*n*) as plotted from left to right: **a-d:** *n* = 29, 31; **e:** *n =* 78, 80, 78; **f**: *n* = 78, 77, 77. B-F: Each dot represents one fly. A, E-F: Each line represents the mean sleep from 16 flies per genotype. All columns are mean ± SEM; asterisks represent P-values *P≤0.05, **P≤0.01, ***P≤0.001, ****P≤0.0001, ns = not significant.

**Extended Data Figure 2:**
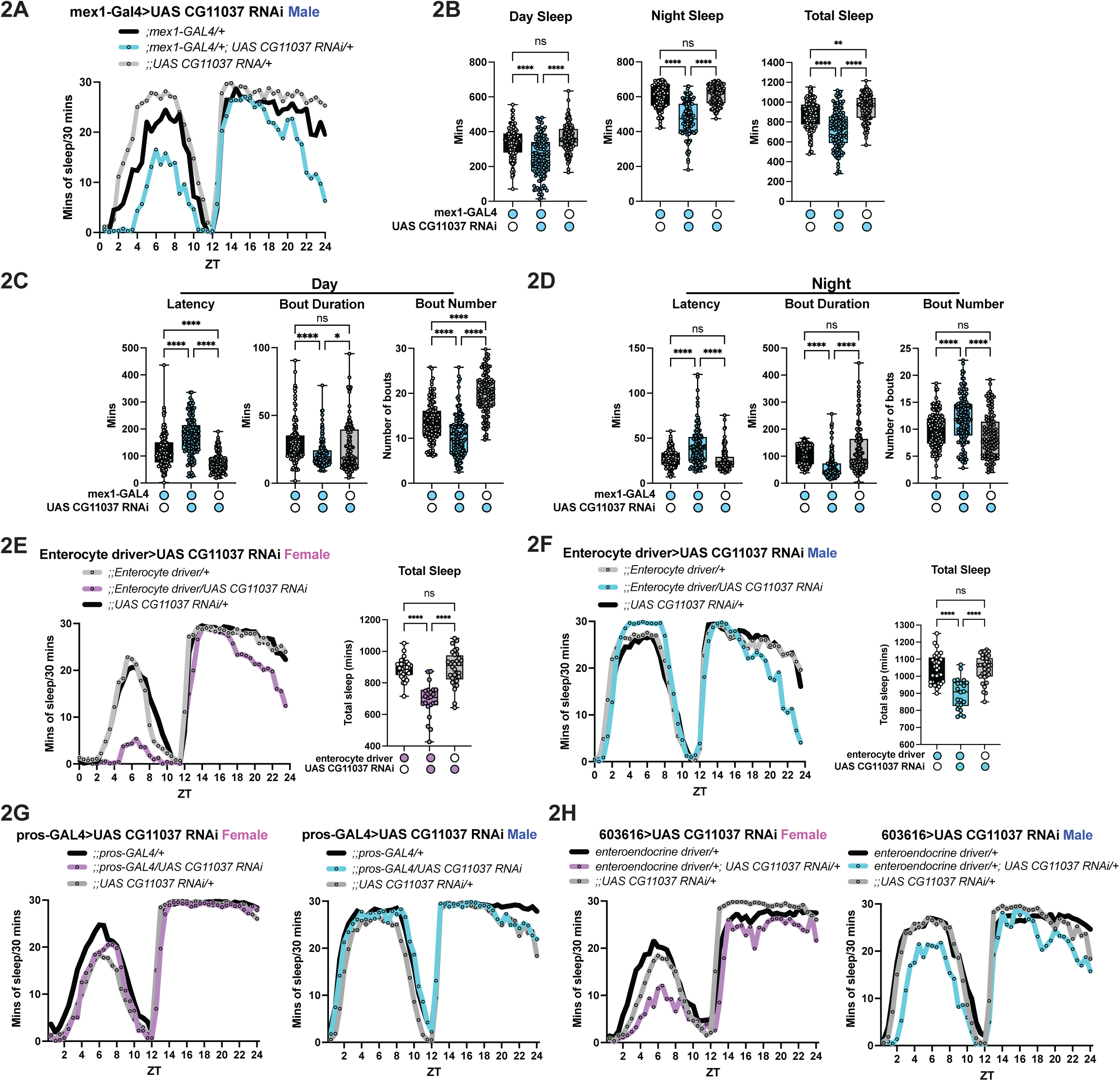
Constitutive expression of *CG11037* is required in enterocytes, not enteroendocrine cells, for daily sleep. (A) Representative sleep trace of male flies with whole life enterocyte knockdown of *CG11037* (blue) compared to *UAS* (gray) and *GAL4* (black) controls. (B) *;mex1-GAL4/+; UAS CG11037 RNAi/+* flies have reduced day sleep (left) and night sleep (middle) leading to overall less sleep (left). (C) Constitutive gut knockdown of *CG11037* increases daytime latency, decreases daytime bout length, and decreases daytime bout number. (D) Constitutive gut knockdown of *CG11037* increases nighttime latency, decreases nighttime bout duration, and increases the overall number of nighttime bouts. (E-F) Knockdown of *CG11037* using an enterocyte split-GAL4 driver (*VT004958-AD; VT004958-DBD*) decreases total sleep in both female (E) and male (F) flies. Sleep traces are quantified on the right. (G) Knockdown of *CG11037* RNAi using *pros-GAL4*, an enteroendocrine cell driver, does not affect female (left) or male (right) sleep. (H) Full life knockdown of *CG11037* using an enteroendocrine split-GAL4 driver (*R20C06-AD; R20C06-DBD)* does not affect female (left) or male (right) sleep. B-F: one way ANOVA with Tukey’s multiple comparisons test. N > 3 biological replicates. The following are the numbers of flies (*n*) as plotted from left to right: **a-d**: *n* = 128, 126, 111; **e**: *n* = 30, 22, 32; **f:** *n =* 32, 22, 32; **g:** *n* = 16, 16, 16, 16, 16, 16; **h:** *n* = 16, 16, 16, 16, 16, 16. B-F: Each dot represents one fly. A, E-H: Each line represents the mean sleep from 16 flies per genotype. All columns are mean ± SEM; asterisks represent P-values *P≤0.05, **P≤0.01, ****P≤0.0001, ns = not significant.

**Extended Data Figure 3:**
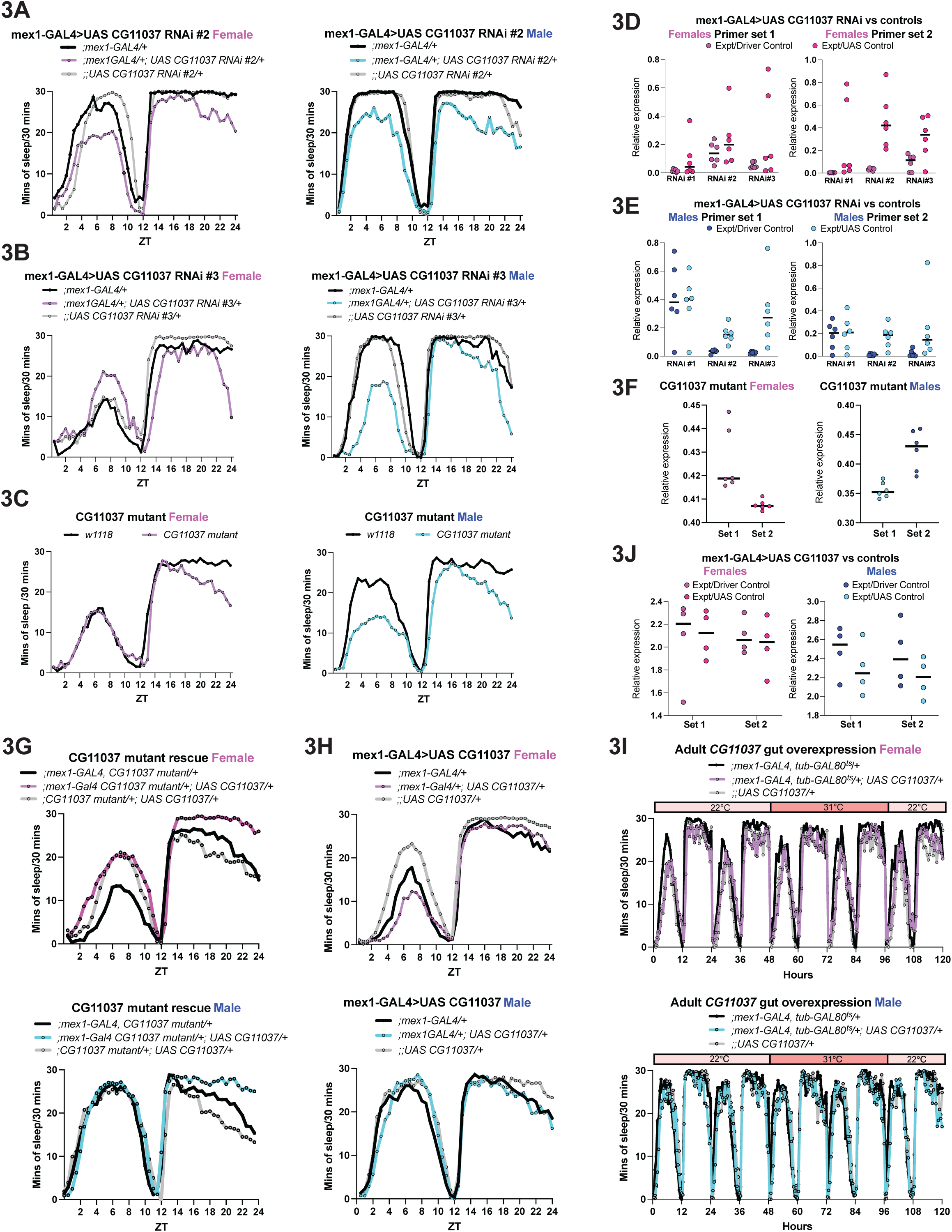
Enterocyte knockdown of *CG11037* decreases sleep, while overexpression has no sleep effect. (A) Constitutive knockdown of *CG11037* using an additional RNAi (RNAi #2) reduces total sleep in females (left) and males (right) with the characteristic decrease in sleep at the end of the night compared to *UAS* and *GAL4* lines. (B) Knockdown of *CG11037* using a third RNAi line decreases sleep in females (left) and males (right) compared to *UAS* and *GAL4* lines. (C) *CG11037* mutants recapitulate decreases in sleep seen using *CG11037* RNAi in both females (left) and males (right) compared to controls. (D-E) Relative RNA expression level measured by qPCR in isolated gut lysates of *;mex1-GAL4/+; UAS CG11037 RNAi* flies compared to *;mex1-GAL4/+* (dark pink, dark blue) or to *;;UAS CG11037 RNAi/+* (light pink, light blue). All RNAis decreased expression, but efficacy varied. (F) Relative RNA expression level measured by qPCR in isolated gut lysates of CG11037 mutants compared to controls. (G) Enterocyte overexpression of *CG11037* rescues *CG11037* mutant sleep compared to controls in *CG11037* mutant backgrounds. (H) Constitutive gut overexpression of *CG11037* in wild-type flies does not increase sleep compared to controls. (I) Adult only overexpression of *CG11037* using the TARGET system *(;mex1-GAL4, tub-GAL80^t^s/+; UAS CG11037/+*) in enterocytes does not increase sleep compared to controls. (J) Relative RNA expression levels measured by qPCR in isolated gut lysates of CG11037 enterocyte overexpression compared to *GAL4* controls (dark pink, dark blue) and *UAS* controls (light pink, light blue). All sleep experiments N > 3 biological replicates, All qPCR experiments N < 4 biological replicates. A-C, G-I: Each line represents the mean sleep of 16 flies per genotype. D-F, J: Each dot in qPCR experiments is one replicate with each replicate encompassing at minimum 40 dissected guts.

**Extended Data Figure 4:**
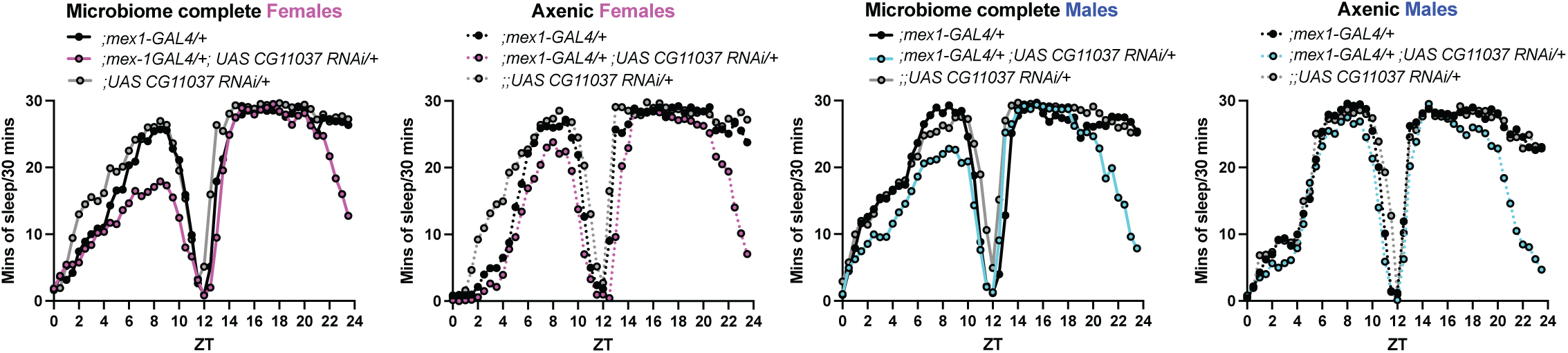
*CG11037* exerts its sleep effects independent of the microbiome. Representative sleep traces of microbiome complete *CG11037* gut knockdown flies (purple or blue solid lines) display the characteristic decrease in sleep at the end of the night compared to *UAS* and *GAL4* controls (gray and black solid lines). Representative sleep traces of axenic *CG11037* gut knockdown flies (purple or blue dotted lines) are still short sleepers compared to *UAS* and *GAL4* controls (gray and black dotted lines). N = 3 biological replicates. All sleep traces represent the mean of 16 flies per genotype.

**Extended Data Figure 5:**
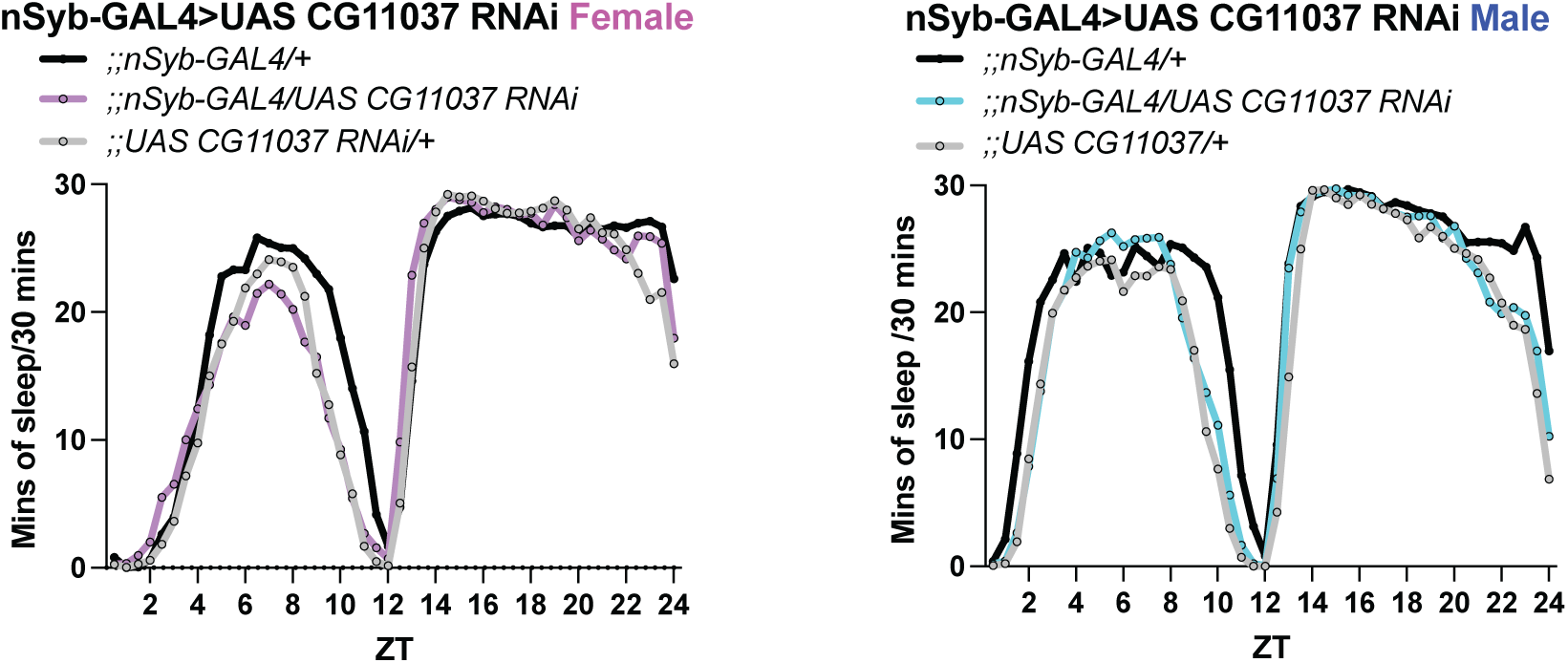
*CG11037* knockdown pan neuronally does not affect sleep. Knockdown of *CG11037* pan neuronally *(;;nSyb-GAL4/UAS CG11037 RNAi*) does not affect daily sleep compared to *UAS* and *GAL4* controls. N > 3 biological replicates. All sleep traces represent the mean of 16 flies per genotype.

**Extended Data Figure 6:**
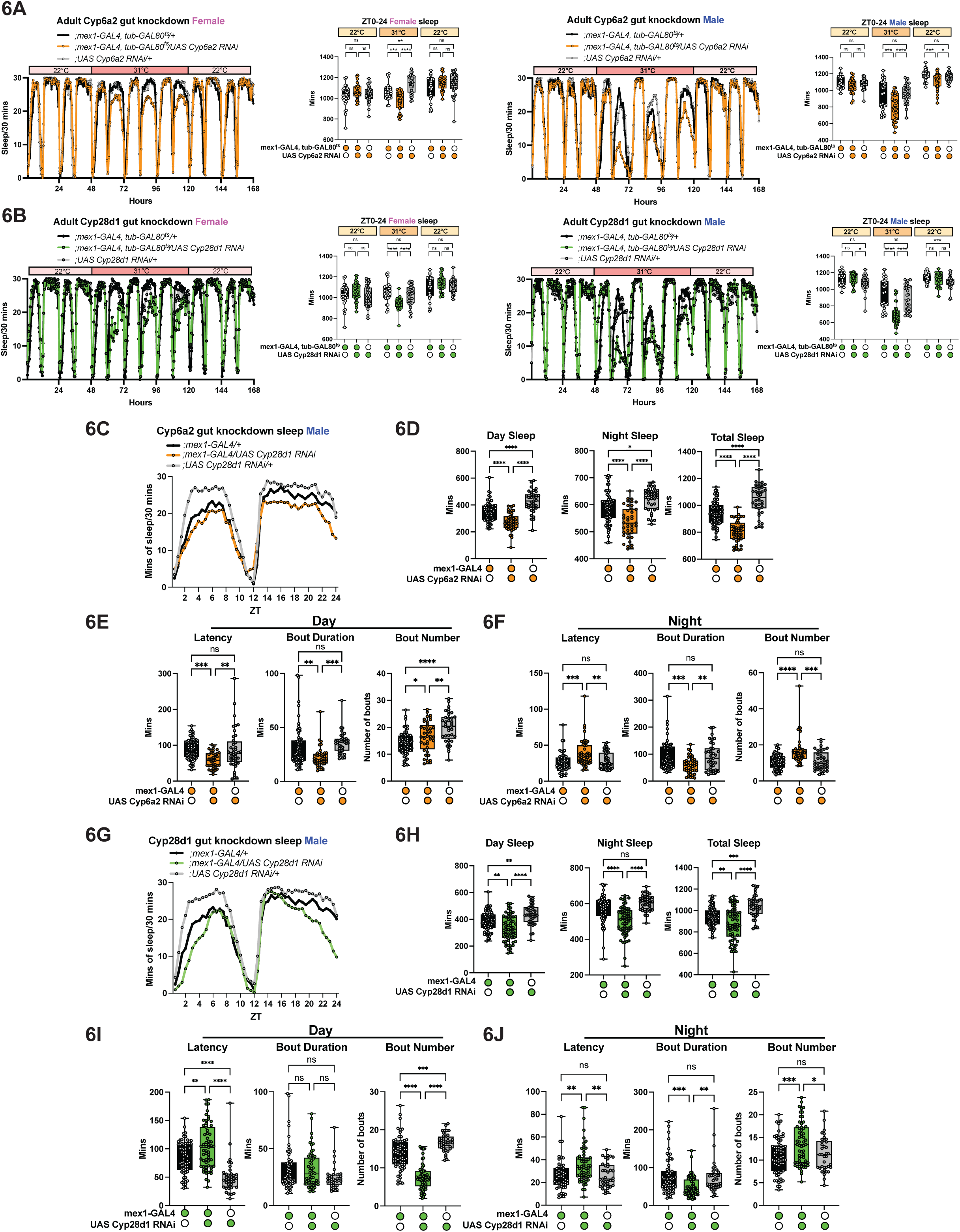
*Cytochrome P450 6a2* and *28d1* are required in enterocytes during adulthood and whole life for daily sleep regulation. (A) Adult restricted knockdown of *Cyp6a2* (orange) in both females (left) and males (right) decreases sleep compared to *UAS* (gray) and *GAL4* (black) controls. Sleep traces are quantified on the right. (B) Adult only knockdown of *Cyp28d1* using the TARGET system (green) decreases sleep in both females (left) and males (right) compared to *UAS* (gray) and *GAL4* (black) controls. Sleep traces are quantified on the right. (C) Representative sleep trace of whole life enterocyte knockdown of *Cyp6a2* (orange) compared to UAS (gray) and GAL4 (black) controls. (D) *;mex1-GAL4/UAS Cyp6a2 RNAi* flies have decreased daytime and nighttime sleep, resulting in an overall decrease in sleep. (E) Constitutive knockdown of *Cyp6a2* decreases daytime latency, decreases daytime bout duration, and has an inconsistent effect on bout number. (F) Enterocyte knockdown of *Cyp6a2* increases nighttime latency, decreases nighttime bout duration, and increases nighttime bout number. (G) Representative sleep trace of *;mex1-GAL4/UAS Cyp28d1 RNAi* (green) compared to *UAS* (gray) and *GAL4* (black) controls. (H) Constitutive enterocyte knockdown of *Cyp28d1* decreases day sleep, night sleep, and total sleep. (I) Enterocyte knockdown of *Cyp28d1* increases daytime latency, does not affect daytime bout duration, and decreases daytime bout number. (J) Enterocyte knockdown of *Cyp28d1* increases nighttime latency, decreases nighttime bout duration, and increases nighttime bout number compared to controls. A-B, D-F, H-J: one way ANOVA with Tukey’s multiple comparisons test. N > 3 biological replicates. The following are the numbers of flies (*n*) as plotted from left to right: **a-b**: *n* = 32, 32, 32; **d-f**: *n* = 64, 42, 47; **h-j:** *n =* 64, 42, 38; A-B, D-F, H-J: Each dot represents one fly. A-B, C, G: Each line represents the mean sleep from 16 flies per genotype. All columns are mean ± SEM; asterisks represent P-values *P≤0.05, **P≤0.01, ***P≤0.001, ****P≤0.0001, ns = not significant.

**Extended Data Figure 7:**
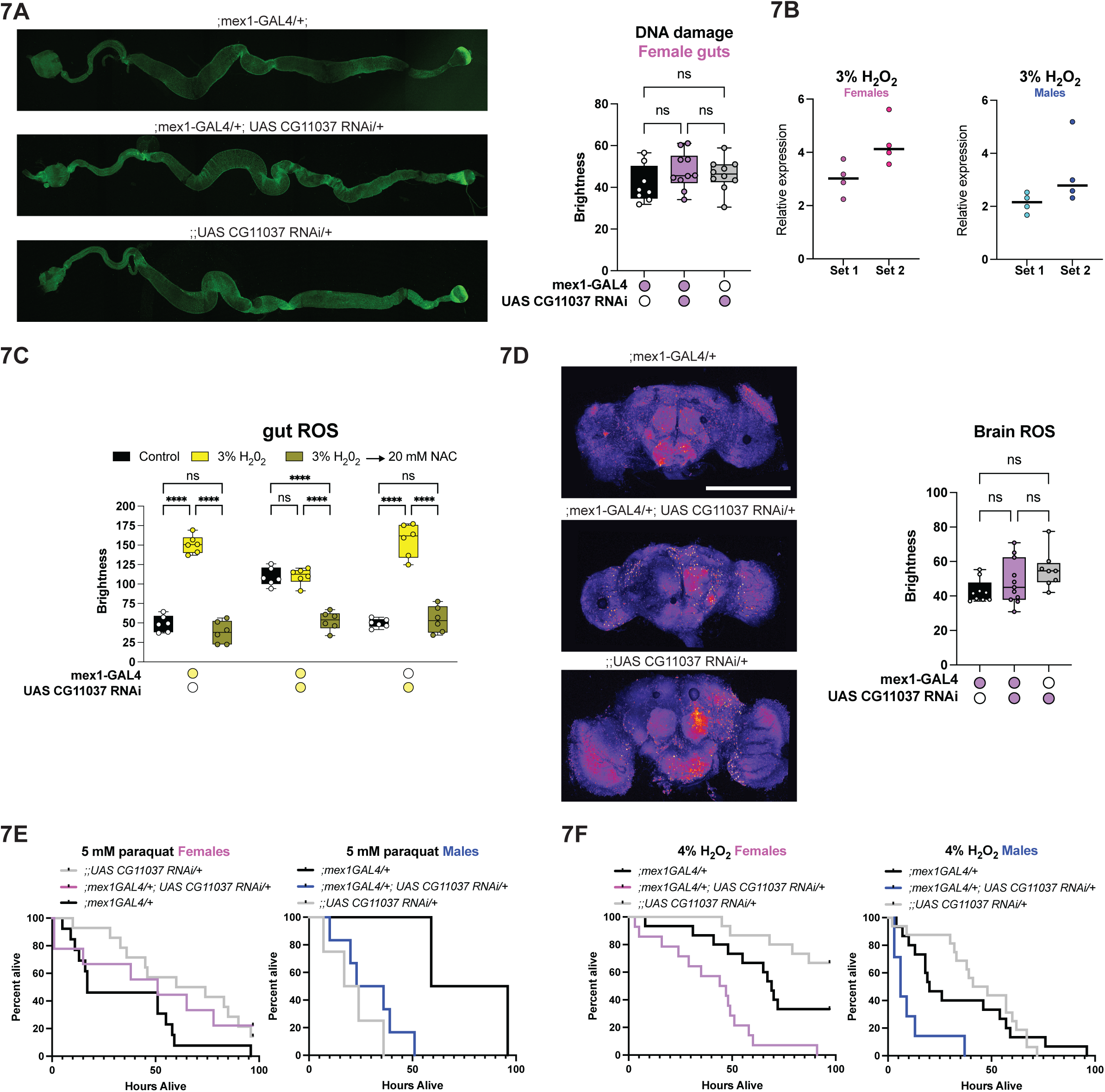
Gut *CG11037* regulates oxidative stress resilience independently of tissue damage. (A) Representative images of tissue damage as detected by H2A staining in *;mex1-GAL4/+; UAS CG11037 RNAi*/*+*, *UAS* and *GAL4* control dissected guts (left). Quantification of H2A staining shows there is no difference in gut DNA damage in enterocyte *CG11037* knockdown flies compared to controls (right). (B) Relative RNA expression levels measured by qPCR in isolated guts from wild-type animals challenged with 3% hydrogen peroxide for 24 hours compared to controls. (C*) ;mex1-GAL4/+; UAS CG11037 RNAi/+* flies have high baseline levels of ROS compared to UAS and GAL4 controls (black). Unlike controls, *;mex1-GAL4/+; UAS CG11037 RNAi/+* flies challenged with 3% hydrogen peroxide for 24 hrs do not increase ROS levels compared to controls (yellow). Subsequent treatment with the antioxidant NAC suppresses accumulation in experimental and control animals (green). (D) Representative images of dissected brains stained with DHE to detect ROS accumulation in *CG11037* enterocyte knockdown flies compared to GAL4 and UAS controls (left). Quantification of ROS accumulation shows there is no difference in brain ROS at baseline (right). (E-F) Survival of female (left) and male (right) flies exposed to 5 mM paraquat (E) or 4% hydrogen peroxide (F). *CG11037* flies are more sensitive to hydrogen peroxide (F) compared to paraquat exposure (E). A, C, D-F: one way ANOVA with Tukey’s multiple comparisons test. E-F: Kaplan-Meier with Log-rank test. N > 3 biological replicates. A, C-D: Each dot represents one fly. Each imaging experiment used at minimum 6 flies. E-F: Survival experiments used 32 flies per genotype per condition. B: Each dot in qPCR experiments is one replicate with each replicate encompassing at minimum 40 dissected guts. All columns are mean ± SEM; asterisks represent P-values ****P≤0.0001, ns = not significant.

**Extended Data Figure 8:**
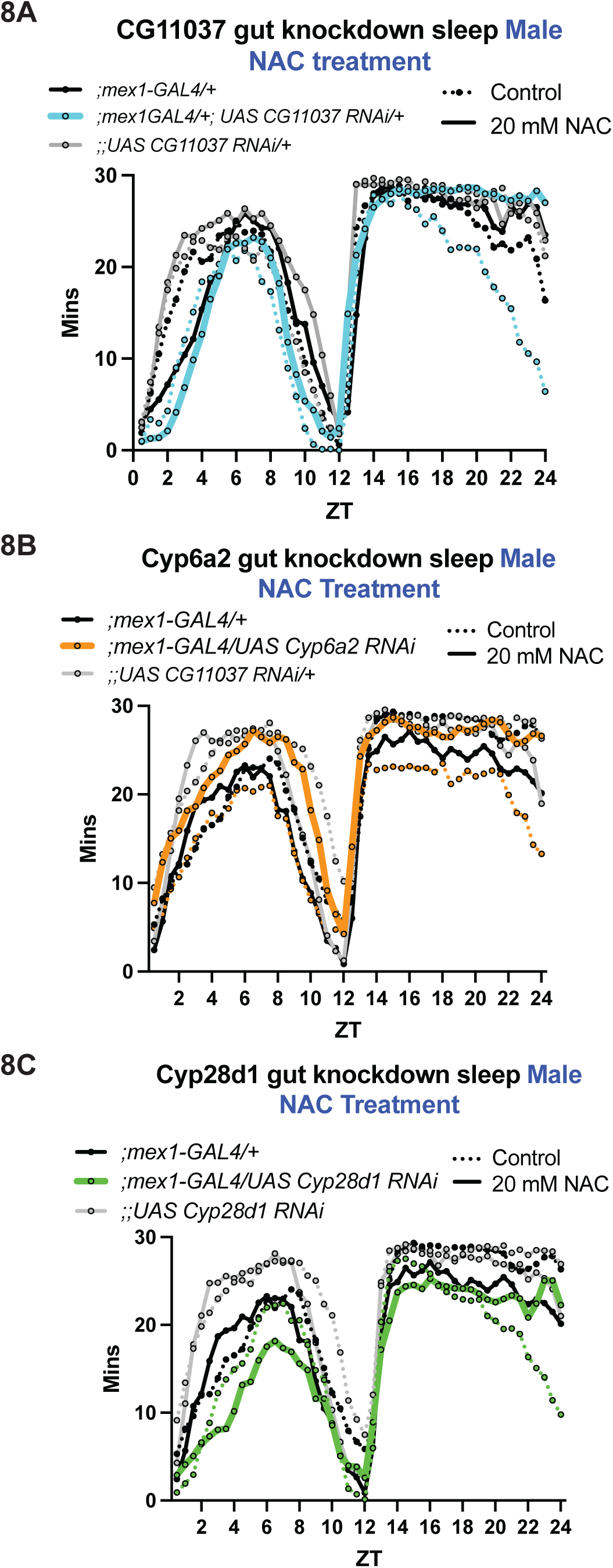
The sleep phenotype of *CG11037* and *Cytochrome P450s* knockdown is ROS dependent. (A) Enterocyte knockdown of *CG11037* reduces total sleep (dotted blue line). Antioxidant treatment with 20 mM NAC restores sleep (solid blue line) back to control levels. Antioxidant treatment has no effect on total sleep in *UAS* (gray lines) or *GAL4* controls (black lines). (B) Whole life gut knockdown or *Cyp6a2* (dotted orange line) decreases daily sleep compared to *UAS* (gray dotted lines) and *GAL4* (black dotted lines) controls. Antioxidant treatment with 20 mM NAC increases *;mex1-GAL4/UAS Cyp6a2 RNAi* (solid orange line) sleep like *UAS* and *GAL4* controls. (C) Enterocyte knockdown of *Cyp28d1* (dotted green line) decreases sleep compared to *UAS* (gray dotted line) and *GAL4* (black dotted lines) controls. Reducing ROS accumulation through antioxidant treatment increases *Cyp28d1* knockdown (green solid line) sleep to control levels (gray and black solid lines). N > 3 biological replicates. Each line represents the mean sleep of 16 flies per genotype per condition.

**Extended Data Figure 9:**
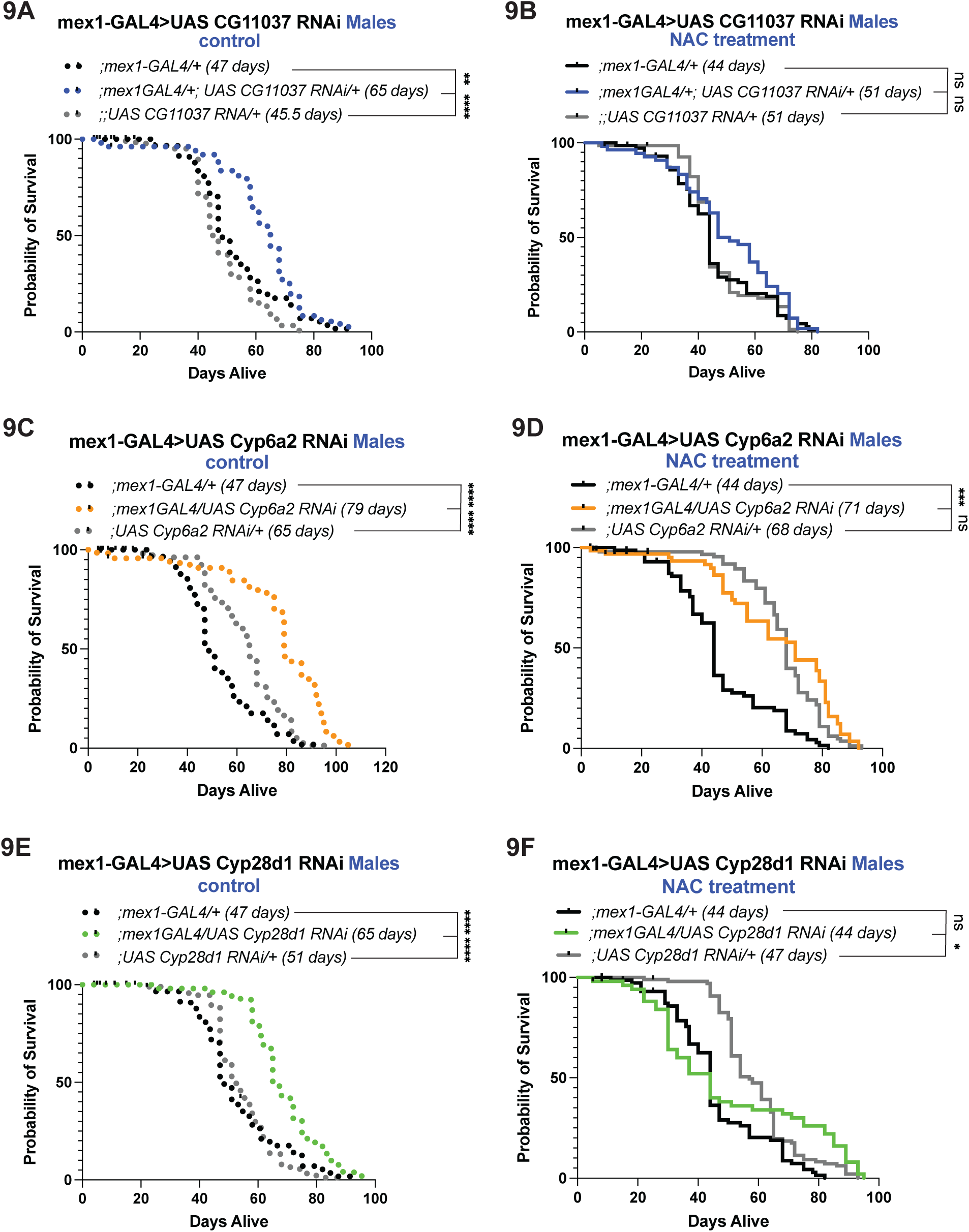
Lifespan extension is ROS dependent. (A, C, E) Whole life knockdown of *CG11037* (A), *Cyp6a2* (C), or *Cyp28d1* (E) extend lifespan compared to *;mex1-GAL4* (dotted black line) and *UAS-RNAi* (dotted gray line) controls. (B, D, F) Whole life treatment with NAC, an antioxidant, decreases lifespan extension in *CG11037* (B), *Cyp6a2* (D), or *Cyp28d1* (F) enterocyte knockdown flies. A-F: Kaplan-Meier with Log-rank test. N > 3 biological replicates. Each survival experiment begins with a minimum of 80 flies.

## Methods

### Fly maintenance and stocks

Flies (*Drosophila melanogaster)* were reared at 25°C and 65% humidity on a 12-hour light:12-hour dark (LD) light cycle and fed a molasses-based food source (Sehgal Lab Standard Yeast-Molasses Diet: 64.7g/L corn meal, 27.1g/L dry yeast, 8g/L agar, 61.6mL/L molasses, 10.2mL/L 20% tegosept, 2.5mL/L propionic acid). *Drosophila* lines used in the RNAi screen originate from the Bloomington Drosophila Stock Center (BDSC, https://bdsc.indiana.edu/) in Indiana, Vienna Drosophila Resouorce Center (VDRC, https://www.viennabiocenter.org/vbcf/vienna-drosophila-resource-center) in Austria, or the Japanese National Institute of Genetics (NIG, (https://shigen.nig.ac.jp/fly/nigfly/) in Japan (Table 1). Gene-switch drivers used for the original sleep screen include gut (TiGS-2, gift from S. Pletcher), fat body (8151), and muscle (43641). Control lines used in the original sleep screen include mCherry RNAi (35785) and LacZ RNAi (60100).

Heavily used fly lines in the manuscript include *w1118* control; RNAi lines UAS CG11037 RNAi #1 (11037R4), UAS Cyp28d1 RNAi (53892), UAS Cyp6a2 RNAi (64008), mex1-GAL4 (91368). All heavily used lines were outcrossed to a *w1118* background 6x. Additional lines used: iso31, nSyb-GAL4, 71993 (Enterocyte driver), 72544 (Enterocyte driver), pros-GAL4 (84276), enteroendocrine driver (603616), CG11037 RNAi #2 (53360), UAS CG11037 RNAi #3 (108803), CG11037 mutant (33153). Mated flies were used for all studies. Sex and age varied by experiment, as specified in method details.

### Genetic screen

All RNAi lines used for the genetic screen can be found in Table 1. All gene-switch drivers used in the genetic screen can be found under fly maintenance and stocks. Flies were raised on standard Sehgal lab food (described above) and maintained at 25°C and 65% humidity on a 12:12 LD cycle. Mated male and female flies aged 4-7 days post eclosion were loaded into locomotor tubes containing 5% sucrose and 2% agar supplemented with 500 μM mifepristone (RU+ food). Control groups were fed food containing equivalent concentrations of 80% ethanol as a vehicle. The TRiP collection of RNAi lines was compared to the UAS-mCherry RNAi line (35785) and the NIG collection of RNAi lines as well as the VDRC RNAi lines are compared to the UAS GFP RNAi line (60100).

Sleep assays were carried out in a 12:12 LD cycle at 25°C and 65% humidity using the Drosophila Activity Monitor System (Trikinetics, http://www.trikinetics.com/). Sleep was analyzed two days after the flies were loaded (∼36 hours after) for 5 days. In initial screening, 8 mated males and 8 mated females per line were tested. Flies were considered asleep when there was no activity for at least five minutes as previously described. Sleep time was analyzed using Insomniac3 as described^1^.

### Sleep recording and analysis

For recording fly sleep, mated 5- to 7-day old flies were loaded into glass tubes containing 5% sucrose with 2% agarose at one end and plugged with cotton/acrylic yarn at the other end for all baseline experiments. For all antioxidant feeding experiments. 5% sucrose with 2% agarose food was supplemented with 20 mM N-Acetyl L-cysteine or supplemented with an equal volume of water.

For all experiments utilizing a gene-switch driver, 5% sucrose with 2% agar food was supplemented with 500 μM RU+ food diluted in 80% ethanol (RU-food).

Briefly, monitors were placed into incubators maintained at 25°C, for GAL4 and gene-switch experiments, in a 12:12 hours light: dark cycle and 50% humidity. For sleep recording using the TARGET system (;mex1-GAL4; tub-GAL80^ts^), baseline sleep was recorded for two days at 22°C, then temperature was raised to 31°C to induce RNAi knockdown for one or two days, before returning to 22°C for recovery sleep.

In these assays, activity counts correspond to breaks of one or more infrared beams that bisect the tube. At least two days after loading flies into single-beam monitors, sleep was analyzed and averaged over 5 days. Sleep was quantified by the *Drosophila* Activity Monitor (DAM) system, using the established definition of a minimum 5 minutes of inactivity^2,3^. Sleep latency was determined from the time of lights-on or from the time of lights-off. Data analysis was performed using Insomniac3 as described^1^.

### Rearing axenic *D. melanogaster*

All protocols for raising axenic flies were adopted from previously published protocols^4^. Briefly, we made axenic flies by surface-sterilizing embryos and growing them in sterile vials. We transferred approximately 15 males and 15 females 3-to 5-day old adult flies to oviposition cages (100 mL plastic beaker, approximately height = 7 cm, diameter = 5.5 cm) capped with a 100 cm Petri dish of sterilized apple juice agar (9g agar, 9g sucrose, 10 mL of 20% Nipagin M in 300 mL of distilled waste and 100 mL of apple juice). We allowed mated females to lay embryos overnight. To sterilize and remove the microbiome from embryos, in a sterilized tissue culture hood we transferred batches of embryos into 40 μm cell strainer, and surface-sterilized them by submerging them in 2 mins in 2.5% active chlorine (50% bleach) followed by 70% ethanol for 2 mins, followed by 2 mins in autoclaved distilled water. The embryos were then transferred onto autoclaved sterile food (Sehgal Lab Standard Yeast-Molasses Diet with antibiotics at final concentrations: 416.7 μg/mL tetracycline, 41.67 μg/mL ampicillin, and 8.333 μg/mL erythromycin). Embryos were left to hatch and develop and were used for experiments at 3-to 5-day old adults.

We confirmed that this method generated axenic flies. From each vial in our experiments, we homogenized 5 mixed sex flies in LB and performed serial dilution in LB (1:10^4^ or 1:10^5^). We plated homogenates on MRS plates (61.15 g/L MRS agar, 1 mL Tween 80, in distilled water), Mannitol plates (1 g/L Beef extract, 10 g/L Peptone, 75 g/L NaCl, 10 g/L Mannitol, .025 g/L Phenol red, Agar 15 g/L, in distilled in water), and LB plates (32 g/L LB Agar, in distilled water). Plates were incubated at 37°C overnight and checked for bacterial growth the next day.

### Generating UAS CG11037 overexpression line

Full length CG11037 cDNA clones were obtained from the *Drosophila* Genomics Resource Center (DGRC) and were used for UAS line creation. cDNA clones were transformed into DH5alpha competent cells and sequenced verified. Both cDNA clones and pUAS-attB were digested using Ecor1 and Xho1 restriction enzymes. Digest products were run out on an agarose gel and cleaned up using a DNA Gel Extraction Qiagen kit. Next, PCR products were ligated together using T4 DNA Ligase and transformed into DH5alpha competent cells. Final plasmids were shipped to BestGene Inc. (http://www.thebestgene.com) for ϕC31 integrase specific germline transformation via embryo infection.

### Mass Spectrometry Sample Preparation

6 biological replicates of bodies and hemolymph were collected for Mass Spectrometry. Siblings from each biological replicate were phenotyped prior to collection.

### Hemolymph collection

Only female flies were collected for these experiments. Groups of 20 5-8-day post-eclosion flies were anesthetized with CO2, rapidly pricked in the thorax with a 27G PrecisionGlide Needle, and loaded into 0.5 mL Eppendorf’s perforated at the base with 22G PrecisionGlide Needle, which were nested inside of a 1.5 mL Eppendorf’s. Nested tubes were then centrifuged for 5000 rpm for 5 min at 4°C. Roughly 20 μL of hemolymph was collected and immediately frozen using dry ice.

### Body collection

Only female flies were collected for these experiments. Groups of 10 5-8-day post-eclosion flies were anesthetized with CO2. Forceps were used to carefully remove fly head, legs, wings, and proboscis. Fly bodies were collected in 1.5 mL Eppendorf tubes and immediately frozen using dry ice.

### Protein Extraction

Hemolymph was solubilized in extraction buffer containing 5% sodium dodecyl sulfate, 50mM TEAB (pH 8.5), and cOmplete EDTA free protease inhibitor cocktail. Samples were centrifuged at 3000g for 10 minutes to clarify lysate.

Fly bodies were solubilized in extraction buffer containing 5% sodium dodecyl sulfate, 50mM TEAB (pH 8.5), and cOmplete EDTA free protease inhibitor cocktail and homogenized using a Preomics Beatbox 96x tissue kit for 20 minutes on high. To shear DNA, samples were sonicated for 10 minutes at 20°C in a Covaris R230 focused-ultrasonicator with the following settings: Dithering: Y=3.0, Speed=20.0, PIP: 360.0, DF: 30, CPB: 200. Samples were centrifuged at 20000g for 10 minutes to clarify lysate.

### In-Solution Digestion

Proteins were processed using the S-Trap protocol^5^, including reduction with TCEP, alkylation with iodoacetamide, and acidification prior to loading onto S-Trap columns. Fly body protein was washed additionally with chloroform:methanol (1:1, v/v). Samples were digested overnight at 37°C with trypsin and LysC. Peptides were then eluted and dried. Samples were then desalted using a 30µm Oasis HLB µElution plate (Waters). Wells were conditioned two times with 200 µL of acetonitrile and equilibrated three times with 200 µL of 0.1% TFA. Samples were applied, washed three times with 200 µL 0.1% TFA, and eluted directly into autosampler vials in three 90µL increments of 50% acetonitrile/0.1% TFA in water. Eluates were then dried by vacuum centrifugation and reconstituted in 0.1% TFA containing iRT peptides (Biognosys, Schlieren, Switzerland). Peptide concentration was determined at OD280 using a Synergy H1 microplate reader (BioTek), and samples were adjusted to 400 ng/ul for injection.

### Mass Spectrometry Data Acquisition

Samples were randomized and analyzed on an Exploris 480 mass spectrometer (Thermofisher Scientific San Jose, CA) coupled with an Ultimate 3000 nano UPLC system and an EasySpray source. 5ul of sample was loaded onto an Acclaim PepMap 100 75um x 2cm trap column (Thermo) at 5uL/min, and separated by reverse phase (RP)-HPLC on a nanocapillary column, 75 μm id × 50cm 2um PepMap RSLC C18 column (Thermo). Mobile phase A consisted of 0.1% formic acid and mobile phase B of 0.1% formic acid/acetonitrile. Peptides were eluted into the mass spectrometer at 300 nL/min with each RP-LC run comprising a 105-minute gradient from 3% B to 45% B. Data independent acquisition (DIA) mass spectrometer settings were as follows: one full MS scan at 120,000 resolution, with a scan range of 350-1200 m/z and normalized automatic gain control (AGC) target of 300%, and automatic maximum inject time. This was followed by variable (DIA) isolation windows, MS2 scans at 30,000 resolution, a normalized AGC target of 1000%, and automatic injection time. The default charge state was 3, the first mass was fixed at 250 m/z, and the normalized collision energy for each window was set at 27.

### Mass Spectrometry QA/QC and System Suitability

The suitability of the instrumentation was monitored using QuiC software (Biognosys; Schlieren, Switzerland) for the analysis of the spiked-in iRT peptides. As a measure for quality control, we injected standard E. coli protein digest in between samples and collected the data in data dependent acquisition (DDA) mode. The collected DDA data were analyzed in MaxQuant^6^ and the output was subsequently visualized using the PTXQC^7^ package to track the quality of the instrumentation.

### Database Searching

The DIA raw files were processed using Spectronaut 19.2 in direct DIA mode^8^. We utilized a Drosophila melanogaster database comprising canonical and reviewed isoforms (23,535 protein entries) from Uniprot, supplemented with a list of 245 common protein contaminants and iRT peptides. Enzyme specificity was set to trypsin with allowance for two potential missed cleavages. Fixed modification was specified as carbamidomethyl of cysteine, while protein N-terminal acetylation and oxidation of methionine were considered variable modifications. To ensure high confidence, a false discovery rate limit of 1% was applied for precursors, peptides, and proteins identification, while the remaining search parameters were maintained at their default settings.

### Bioinformatic Processing

Proteomics data processing and statistical analysis were performed in R using MS2 intensity values generated by Spectronaut. Data were log2-transformed and normalized by median-centering each sample. Proteins were retained for downstream analysis only if detected in at least 5 out of 6 replicates within at least one experimental cohort.

Differential abundance analysis was performed using the DEP package in R with Limma moderated t-tests. Upon reevaluation of the dataset, the two control cohorts were treated as distinct genotypes rather than combined into a single control condition. Differential expression analyses were therefore conducted by separately comparing the mutant cohort against each control genotype using the processed and normalized intensity dataset provided by the core facility.

Volcano plots were generated to visualize differential protein abundance and statistical significance for each comparison. The y-axis represents the −log10 adjusted P value, and proteins with an adjusted P value < 0.25 were considered significantly altered and prioritized for downstream bioinformatics analyses.

Data is available using Pride: PXD079043.

### Lifespan Experiments

For all lifespan studies, virgin male and female flies were collected under CO2 anesthesia and allowed to mate for 48 hours on standard food at a maximum density of 30/vial (15 males and 15 females) Following 48 hours of mating, male and female flies were separated by sex and housed single-sex on standard food or antioxidant supplemented food (20 mM NAC) at a maximum density of 25/vial with a minimum of 80 flies per genotype and experiment, repeated a minimum of three times. For all lifespan studies flies were flipped (no CO2) to fresh vials and dead/escaped were tallied every 3-4 days. Each vial was longitudinally maintained on the same assigned food condition and survival was tracked until all flies were dead. Flies were never again anesthetized after lifespan monitoring began.

Food: For all lifespan studies, fresh food was made no more than 1 week prior to use and maintained at 4°C prior to use. Pre-made yeast-molasses food was melted in the microwave and supplemented with 20 mM NAC diluted in water or MilliQ water. 2 mL of control or NAC supplemented food was poured on top of standard lab food to be used for lifespan studies. Food vials were warmed to room temperature prior to use.

### Sleep deprivation

To sleep deprive flies, we used mechanical sleep deprivation. To achieve deprivation, flies in up to four single beam Trikinetics monitors were fixed to a multi-tube vortex machine (VWR) filled with a mounting plate. The vortexer was attached to an LC4 light controller and programmed to be shaken randomly for 2s every 20s over a 12-hour period (ZT12-ZT24) for one night. To estimate rebound sleep, 4 or 12 hours of sleep after sleep deprivation as compared with sleep on the pre-deprivation day at the same ZT time (ZT0-ZT4, or ZT0-ZT12). Sleep latency was measured using Insomniac3 software^1^.

### qPCR

Adult fly guts (∼40) were dissected in PBS and subjected to TRIzol extraction (Invitrogen Cat# 15596026). 500 μg of RNA was reverse transcribed to generate cDNA using a high-Capacity cDNA Reverse Transcription kit (Invitrogen Cat# 4368814). qPCR was performed on a QuantStudio 7 Pro Real-Time PCR system (Thermo Fisher Scientific) using SYBR Green PCR master mix (Applied Biosystems Cat# 4309155). Relative expression was determined using the ΔΔCT method. For each sample, mean CT values were determined from 3 technical replicates. ΔCT was determined relative to the house keeping gene, *α-Tubulin*. ΔΔCT was then calculated as fold change relative to the control group. Real-time qPCR primers were sourced from the FlyPrimerBank or previous publications. Primer sequences can be found in Table S4.

Oxidative stress exposed flies: Day 5-7 post eclosion wild type (w1118) mated male and female flies were exposed to 3% hydrogen peroxide (supplemented in complete media) for 24 hrs. Following 24 hrs of hydrogen peroxide exposure guts were dissected in PBS and collected in TRIzol for RNA extraction. Primers were either designed de novo using https://www.idtdna.com or http://flyrnai.org/flyprimerbank or from citations listed.

### Immunostaining

Guts were dissected in PBS and fixed in 4% paraformaldehyde in PSB for 1 hr rocking at room temperature. Guts were then rinsed three times in PBS + 0.1% Triton X-100 (Sigma Aldrich, Cat# 9036-19-2), permeabilized, and blocked in 2% bovine serum albumin (BSA) PBT for 1 hr at room temperature. Samples were then incubated with Anti-Histone H2AvD pS137 (1:500, Thomas Scientific Cat# 600-401-914S) diluted in blocking serum overnight rocking at 4°C. Samples were then washed three times in PBT and Goat-anti Rabbit Alexa Fluor 488 (1:1000, Life technologies Cat# A32731) was applied at room temperature rocking for 2 hr. Samples were further washed four times in PBT and mounted in vectashield antifade mounting medium (Vector Laboratories Cat# H-1000-10). All images were obtained with a Leica Stellaris STED confocal microscope and processed using FIJI.

### DHE staining and quantification

For ROS measurement with Dihydroethidium (DHE, Invitrogen Cat# D11347) stock solutions of 10 mM DHE in DMSO (Life technologies Cat# D12345) were made. Guts or brains were dissected in Schneider’s insect medium (Life technologies Cat# 21720024). Immediately following dissection guts/brains were incubated in 60 μM DHE diluted in Schneider’s insect medium for 7 mins rocking at room temperature covered with aluminum foil to shield from light exposure. Guts/brains were then washed twice (5 mins) in Scheider’s insect medium, and then once in PBS at room temperature, rocking and covered with aluminum foil.

Finally, guts/brains were mounted and immediately imaged every 3 μm using a Lecia Stellaris STED confocal microscope without fixation. Gut images were acquired by 10x (dry), and brains by 20x (dry) objectives. To ensure that guts were not squished, we lined each end of the slide with three layers of double-sided tape prior to placing a cover slip down. Key point: Flies were anesthetized using ice, not CO2. Our experiences suggests that CO2 can change DHE staining results.

ROS exposure: For experiments in which ROS was measured following oxidative stress. A subset of siblings were used to measure baseline ROS as described above. Leftover siblings were placed in vials containing complete food supplemented with 3% hydrogen peroxide for 24 hours and then ROS was measured as described above. Finally, remaining siblings are moved into vials containing complete food supplemented with 20 mM NAC (diluted in water) for 24 hours and then ROS was measured as described above.

### Sickness sleep and immune response measurements

All methods used for behavioral assays and for measuring immune function are described in detail elsewhere^16^. All sickness sleep and immune response measurements utilized 5-7 day old mated female flies. Briefly, *S. marcescans* (ATCC, Cat# 8100) were grown overnight in Luria Broth (LB) medium and diluted to an OD_600_ of 0.05 in Phosphate Buffered Saline (PBS) with 1% food coloring (Amazon Cat# B0CJ2L78SR). Flies were loaded into Drosophila Activity Monitors for three days to acclimate and record baseline sleep. On day 3 monitors were removed at ZT18, flies were CO2 anesthetized, and infected by injection with diluted bacteria using glass capillary needles or subjected to aseptic injury by injection with equivalent dilutions of PBS with 1% food coloring. Handled control flies were similarly removed from monitors, anesthetized on CO2 for a similar amount of time septic/sterile injury took, and then returned to their monitors. Flies were immediately returned to incubators until maintained in incubators until all septic injury flies died. Survival rate was determined by using activity data derived from the Trikinetics system data.

Sickness Sleep Response: Sickness sleep was measured as previously described^17^. In brief, sickness sleep was calculated in individual flies as the difference in sleep (minutes/4 hrs) between the day after infection/injury and the equivalent time interval the day before injury. These differences were adjusted by subtracting the average difference for corresponding timepoints from handled controls animals.

Post-infection sleep was measured and reported only in flies that survived beyond the analysis period of 24 hours.

Bacterial Load: To determine bacterial load, groups of 10 infected flies were homogenized in 400 μL LB media immediately or 24 hrs post-injection with bacteria. Homogenized samples were serially diluted (1:10^2^ or 1:10^3^ for flies homogenized immediately after infection, 1:10^4^ or 1:10^5^ for flies homogenized 24 hrs post-infection) and spread onto LB agar plates. LB plates were incubated at 37°C overnight, and the number of colony forming units (CFUs) per fly was determined via manual count. CFUs per fly were determined by normalizing the number of colonies counted on LB plates to the dilution factor, plated volume, number of flies pooled per samples, and the initial homogenate volume.

Survival: Survival rate was determined by using activity data derived from the Trikinetics monitoring system. Survival was measured in hours following infection. Flies were considered dead when all activity counts reached zero for the remainder of the experiment.

### Oxidative stress survival experiments

Day 5-7 post-eclosion mated male and female flies were transferred to individual Drosophila Activity Monitor tubes containing 5% sucrose and 2% agar supplemented with 5 mM paraquat (Sigma Aldrich Cat# 856177), 4% hydrogen peroxide (Fisher Scientific Cat# BP2633500), or water. Survival rate was determined by using activity data derived from the Trikinetics monitoring system. Survival was measured immediately from the moment flies encountered oxidative stress and measured until flies were all dead (i.e. activity counts reached zero for the remainder of the experiment).

### Statistics

For all experiments, except those related to mass spectrometry, Prism was used to conduct all statistical testing and produce plots. All quantifications are displayed with box plots where the box surrounds points within the interquartile range, a line within the box representing the median, and bars extending to total range of data points, except where noted. All quantifications of 2 groups were analyzed using unpaired t-tests, comparing one group to a single control. All quantifications including 3 groups were analyzed using a one way ANOVA with Tukey’s multiple comparison test. Survival and lifespan experiments were analyzed using log-rank Kaplan Meier tests.

### Key resources table

| REAGENT or RESOURCE | SOURCE | IDENTIFIER |
| --- | --- | --- |
| **Antibodies** |  |  |
| Anti-Histone H2AvD pS137 | Thomas Scientific | Cat# 600-401-914S |
| Goat anti-Rabbit IgG (H+L) Alexa Fluor 488 | Life Technologies | Cat# A32731 |
| **Bacterial and virus strains** |  |  |
| *S. marcescens* | ATCC | ATCC# 8100 |
| **Chemicals, peptides, and recombinant proteins** |  |  |
| 30% Hydrogen Peroxide | Fisher Scientific | Cat# BP2633500 |
| Paraquat | Sigma Aldrich | Cat# 856177 |
| Food coloring | Amazon | Cat# B0CJ2L78SR |
| Dihydroethidium (DHE) | Invitrogen | Cat# D11347 |
| DMSO anhydrous | Life Technologies | Cat# D12345 |
| DPBS | Fisher Scientific | Cat# 14190235 |
| Triton X-100 | Sigma Aldrich | Cat# 9036-19-2 |
| Vectashield Antifade | Vector Laboratories | Cat# H-1000-10 |
| Paraformaldehyde | Electron Microscope Services | Cat# 15710-S |
| Schneider's Insect Media | Life Technologies | Cat# 21720024 |
| N-acetyl L-cysteine | Sigma Aldrich | Cat# A7250 |
| Mifepristone (RU486) | Sigma Aldrich | Cat# M8046 |
| Agar, Drosophila Type | Fisher Scientific | Cat# NC1429200 |
| Corn Meal | LabScientific | Cat# FLY-8010-20 |
| Drosophila Dry Active Yeast | LabScientific | Cat# FLY-8040-10 |
| Molasses | Genesee Scientific | Cat# 62-118 |
| SYBR Green PCR Master Mix | Applied Biosciences | Cat# 4309155 |
| TRIzol | Invitrogen | Cat# 15596026 |
| Nipagin M | Sigma Aldrich | Cat# M8911-1VL |
| MRS agar | Thermo Fisher | Cat# CM0361B |
| Mannitol Salt Agar | Thermo Fisher | Cat# CM0085B |
| Beef extract | Thermo Fisher | Cat# 212303 |
| Peptone | Fisher Scientific | Cat# BP9725-500 |
| Sodium Chloride | Fisher Scientific | Cat# S271-500 |
| Phenol red | Thermo Scientific | Cat# B21710.09 |
| LB Agar | Fisher Scientific | Cat# BP9724-500 |
| Tween 80 | Fisher Scientific | Cat# BP338500 |
| Ampicillin Sodium Salt | Fisher Scientific | Cat# BP1760-25 |
| Tetracycline | Sigma Aldrich | Cat# T3258 |
| Erythromycin | Sigma Aldrich | Cat# E5389 |
| Sucrose | Life Technologies | Cat# A15583.0E |
| Propionic acid Sodium Salt | Genesee Scientific | Cat# 20-271 |
| Cell strainer | Fisher Scientific | Cat# 07-201-430 |
| Ecor1 (fast digest) | Life Technologies | Cat# FD0274 |
| Xho1 (fast digest) | Life Technologies | Cat# FD0694 |
| Sodium Dodecyl Sulfate | Thermo Scientific | Cat# J18220.A7 |
| Triethylammonium bicarbonate buffer (TEAB) | Sigma Aldrich | Cat# T7408 |
| cOmplete, EDTA -free protease inhibitor cocktail | Millipore Sigma | Cat# 4693132001 |
| **Critical commercial assays** |  |  |
| High-Capacity cDNA Reverse Transcription Kit | Invitrogen | Cat# 4368814 |
| QIAquick PCR purification kit | Qiagen | Cat# 28104 |
| T4 DNA Ligase | New England Biolabs | Cat# MS0202S |
| **Deposited data** |  |  |
| Raw and analyzed data | This paper | PRIDE: PXD079043 |
| **Experimental models: Cell lines** |  |  |
| *E. coli*: DH5alpha Competent Cells | Thermo Fisher | Cat# 18258012 |
| **Experimental models: Organisms/strains** |  |  |
| RNAi lines used in screen (stock center backgrounds) | Table S1 | Table S1 |
| *D. melanogaster*: Iso31 control strain | Lab generated allele | N/A |
| *D. melanogaster*: w1118 control strain | Lab generated allele | N/A |
| *D. melanogaster*: nSyb-GAL4 | Lab generated allele | N/A |
| *D. melanogaster*: Enterocyte AD | Bloomington<br>Drosophila Stock<br>Center | BL# 71993 |
| *D. melanogaster*: Enterocyte DBD | Bloomington<br>Drosophila Stock<br>Center | BL# 72544 |
| *D. melanogaster*: Mex1-GAL4 | Bloomington<br>Drosophila Stock<br>Center | BL# 91368 |
| *D. melanogaster*: w1118; R20C06-p65.AD; R20C06-GAL4.DBD | Bloomington<br>Drosophila Stock<br>Center | BL# 603616 |
| *D. melanogaster*: Pros-GAL4 | Bloomington<br>Drosophila Stock<br>Center | BL# 84276 |
| *D. melanogaster*: UAS CG11037 RNAi (RNAi #1) | Fly Stocks of National<br>Institute of Genetics | NIG# 11037R-4 |
| *D. melanogaster*: UAS CG11037 RNAi (RNAi #2) | Bloomington<br>Drosophila Stock<br>Center | BL# 53360 |
| *D. melanogaster*: UAS CG11037 RNAi (RNAi #3) | Vienna Drosophila<br>Resource Center | VDRC #108803 |
| *D. melanogaster*: UAS Cyp28d1 RNAi | Bloomington<br>Drosophila Stock<br>Center | BL# 53892 |
| *D. melanogaster*: UAS Cyp6a2 RNAi | Bloomington<br>Drosophila Stock<br>Center | BL# 64008 |
| *D. melanogaster*: CG11037 mutant | Bloomington Drosophila Stock Center | BL# 33153 |
| *D. melanogaster*: ;mex1-GAL4; tub-GAL80<sup>ts</sup> | This paper | N/A |
| *D. melanogaster*: Digestive tract Geneswitch (TiGS-2) | Gift from S. Pletcher | N/A |
| *D. melanogaster*: at body Geneswitch (S106) | Bloomington Drosophila Stock Center | BL# 8151 |
| *D. melanogaster*: Muscle Geneswitch (MHC) | Bloomington Drosophila Stock Center | BL# 43641 |
| *D. melanogaster*: UAS mCherry RNAi | Bloomington Drosophila Stock Center | BL# 35787 |
| *D. melanogaster*: VDRC GFP control | Vienna Drosophila Resource Center | VDRC# 60100 |
| **Oligonucleotides** |  |  |
| Oligonucleotides used for qPCR | Table S4 | Table S4 |
| **Recombinant DNA** |  |  |
| CG11037 cDNA | Drosophila Genome Resource Center | Cat# AT20289 |
| pUAST-attB | Drosophila Genome Resource Center | Cat# 1419 |
| **Software and algorithms** |  |  |
| GraphPad Prism | GraphPad | Download at http://graphpad.com |
| DAM FileScan | Trikinetics | Free download at http://trikinetics.com |
| Insomniac3 | Lab generated software |  |
| Benchling (plasmid design) | Benchling | http://benchling.com |
| **Other** |  |  |
| R230 Focused-ultrasonicator | Covaris | Cat# 500620 |
| PreOmics Beatbox 96x tissue kit | PreOmics | Cat# P.O.00121 |
| Single-Beam Sleep Monitors | Trikinetics | DAM2 |
| Transgenic Flies | BestGene Inc. | http://www.thebestgene.com |
| QuantStudio 7 Pro Real-Time PCR system | Thermo Scientific |  |
| Forceps Dumont #5SF | Fine Science Tools | Cat# 11252-20 |
| 27G PrecisionGlide Needle | Becton Dickinson | Cat# 305126 |
| 22G PrecisionGlide Needle | Becton Dickinson | Cat# 305155 |

