## Supplementary material for "Gut-derived serine protease reduces sleep but extends lifespan": Table S4

| Oligonucleotides | | |
| --- | --- | --- |
| CG11037 Set 1 Forward  GCTGGCTTGCACTTTAGTTTC | This paper | N/A |
| CG11037 Set 1 Reverse GCCACATCGAGAGTGAGATTT | This paper | N/A |
| CG11037 Set 2 Forward  CGAGAGGACGACATGAATATGG | This paper | N/A |
| CG11037 Set 2 Reverse  TAAGGCTCACCGAACATAAGC | This paper | N/A |
| Relish Forward  AGTGGTCCAAGAAGACAGAAAG | Swanson et al. | N/A |
| Relish Reverse  GAACAGAGCCGGTCGTAAAT | Swanson et al. | N/A |
| Pirk Forward  CGACAGAGACGGAGATAGAGATAG | This paper | N/A |
| Pirk Reverse  CCGCTGCTAATCACTCGTAAA | This paper | N/A |
| Dif Forward  CAGTTTGCTACGACCGGAGAGCTA | Swanson et al. | N/A |
| Dif Reverse  GAATATCCGCCAGTTGCAGAGTGC | Swanson et al. | N/A |
| Dorsal Forward  CAACCCTTTGGGCTTTCTTATC | Swanson et al. | N/A |
| Dorsal Reverse  GTGCTGTTGTGGTTGTAGTTG | Swanson et al. | N/A |
| Cactus Forward  CTGCTCAACATCCAGAACGA | <http://www.flyrnai.org/flyprimerbank> | N/A |
| Cactus Reverse  GCCGAACTTCTCTGTCAAGG | <http://www.flyrnai.org/flyprimerbank> | N/A |
| Upd1 Forward  AGACAGCCGTCAACCAGAC | Moskalev et al. | N/A |
| Upd1 Reverse  GCTTCAAACGCTTGTTCATC | Moskalev et al. | N/A |
| Upd2 Forward  CGGAACATCACGATGAGCGAAT | Rajan and Perrimon. | N/A |
| Upd2 Reverse  TCGGCAGGAACTTGTACTCG | Rajan and Perrimon. | N/A |
| Upd3 Forward  ACTGGGAGAACACCTGCAAT | Woodcock et al. | N/A |
| Upd3 Reverse  GCCCGTTTGGTTCTGTAGAT | Woodcock et al. | N/A |
| STAT92E Forward  CTGGGCATTCACAACAATCCAC | Rajan and Perrimon. | N/A |
| STAT92E Reverse  GTATTGCGCGTAACGAACCG | Rajan and Perrimon. | N/A |
| Wg Forward  CGCAGAGTCGGACGAAAGAA | Suresh et al. | N/A |
| Wg Reverse  TTTCGAGCGGAGGAGTGAAG | Suresh et al. | N/A |
| α-Tubulin Forward  CGTCTGGACCACAAGTTCGA | Xu et al. | N/A |
| α-Tubulin Reverse  CCTCCATACCCTCACCAACGT | Xu et al. | N/A |
